# A conserved molecular marker for connections between two evolutionarily distinct visual centers

**DOI:** 10.64898/2026.08.17.745220

**Authors:** Nicholas M. Oliver, Maia Jin Classe, Sebastian Werneburg, Elise L. Savier

## Abstract

Sensory systems share common circuit organization motifs across mammalian species, however, anatomical subdivisions show varying degrees of complexity depending on ecological niche and species-specific sensory requirements. While coarse neuroanatomical connections seem preserved within the visual system, it remains unknown if molecularly defined cell-types share a similar degree of conservation, regarding not only their functional properties but also connectivity. Here we analyze the organization, molecular marker expression, and connections between two prominent visual centers, the superior colliculus (SC) and the dorsal lateral geniculate nucleus of the thalamus (dLGN), in the mouse and the tree shrew, a highly visual, diurnal species closely related to primates.

Previous attempts to link molecular markers to subdivisions and connectivity of the dLGN have shown lack of conservation across species, thus preventing the systematic investigation of brain-wide interactions involved in vision. Leveraging recent single-cell and single-nucleus RNA sequencing studies, our results unravel a conserved molecular marker that shows spatial restriction in the dLGN and correlates with the location of connections from the SC in both the mouse and the tree shrew. We extend our findings by confirming the presence of this molecular marker in the human dLGN. These results provide a molecular definition and genetic access point for SC to dLGN connections in the mouse and tree shrew, enabling cell-type specific studies of the parallel processing of visual information.

**Significance Statement:** While coarse neuroanatomical connections are conserved across mammalian species, their local projection pattern can vary depending on sensory specialization. Furthermore, it remains unknown if these connections are made by homologous molecularly defined cell-types. Our work confirms the conservation of a projection between the superior colliculus and the visual thalamus in the mouse and the tree shrew, which have distinct visual capabilities. We also uncover a genetic access point, unraveling new avenues to understand how these two major visual centers, often studied separately, interact with each other. The remarkable conservation of a molecular marker suggests that despite core anatomical variations and organization principles, the expression of certain genes linked to brain-wide connectivity can be conserved across species.

## Introduction

Investigating the conservation of neuronal circuits across species provides important insight into how the processing of sensory information is implemented (Tosches, 2021). In the mammalian visual system, the dorsal lateral geniculate nucleus (dLGN) of the thalamus is the first central relay for conveying visual information from the retina to the primary visual cortex (Cruz-Martín et al., 2014; Guido, 2018; Liang and Chen, 2020). In parallel, visual information is also projected from the retina to the superior colliculus (SC, also called tectum in non-mammalian species), a conserved midbrain structure involved in multisensory integration (May, 2006; Cang et al., 2018; Basso et al., 2021). Projections between the SC and dLGN (tectogeniculate) have been found across 19 mammalian species (Harting et al., 1991; Hendry and Yoshioka, 1994; Hendry and Reid, 2000; Merkulyeva, 2022). The evolutionary conservation of the tectogeniculate projection suggests an important functional role, however, our understanding of its contribution to vision remains elusive due to lack of genetic access and differences in anatomical layout across species.

In primates, the dLGN is laminated, and primarily organized around the magnocellular (M) and parvocellular (P) pathways (Shapley and Hugh Perry, 1986; Merigan and Maunsell, 1993; Hendry and Reid, 2000; Rathbun and Usrey, 2009). In addition to these two primary pathways, there is evidence for a third pathway known as the koniocellular (K) pathway which receives tectogeniculate projections (Lachica and Casagrande, 1993; Casagrande, 1994; Hendry and Reid, 2000; Martin and Solomon, 2019). Evidence for heterogeneity in the tectogeniculate projection has been found across species, both morphologically and functionally (Diamond et al., 1991; Gale and Murphy, 2014; Bickford et al., 2015; Fei et al., 2025; Sciaccotta et al., 2025). Further investigation of the cell-types that comprise this projection (tectogeniculate neurons) and neurons that directly receive these projections (tectorecipient neurons) requires the identification of genetic access points. With its genetic tractability, the mouse is a powerful model to investigate neuronal circuitry at a cell-type specific level. However, the mouse dLGN lacks lamination and is organized in two parts: the “core” and the “shell”, the latter receiving tectogeniculate projections (Harting et al., 1991; Grubb and Thompson, 2004; Cruz-Martín et al., 2014; Bickford et al., 2015). Tree shrews (*Tupaia belangeri*) are closely related to primates and possess similar visual system organization (Petry and Bickford, 2019). The tree shrew dLGN is organized into 6 layers separated by interlaminar zones (ILZs) and tectogeniculate projections target layers 3, 6, and an ILZ between layers 4 and 5 (Diamond et al., 1991; Harting et al., 1991; Hendry and Reid, 2000; Petry and Bickford, 2019; Sciaccotta et al., 2025), making the tree shrew an ideal model to study brain-wide interactions involved in visual processing.

Previously identified molecular markers have provided important insights into the subdivisions of the primate dLGN. The M and P layers of the primate dLGN express the calcium binding protein parvalbumin (PV) (Goodchild and Martin, 1998; Ma et al., 2023), and immunostaining of calbindin (CB) enabled the first studies of the K layers (Jones and Hendry, 1989; Diamond et al., 1993; Hendry and Yoshioka, 1994; Johnson and Casagrande, 1995; Goodchild and Martin, 1998; Hendry and Reid, 2000). Recently, spatial transcriptomics and RNA sequencing (RNAseq) have allowed the identification of cell-types and their associated molecular markers (Murray et al., 2008; Tasic et al., 2018; Bakken et al., 2021). Here, by leveraging these RNAseq studies (Bakken et al., 2021; Liu et al., 2023, 2025), we confirm spatial restriction of a previously identified molecular marker, Necab1, to the shell of the dLGN in mice (Bakken et al., 2021) and uncover specificity for tectorecipient layers in tree shrews. We then confirm expression of Necab1 in the human dLGN. Furthermore, using transsynaptic tracing, we show that tectorecipient neurons in the mouse express Necab1 and are mostly glutamatergic. Together, these results provide genetic access to subdivisions of the dLGN that integrate multiple visual streams, furthering our understanding of brain-wide interactions in vision.

## Materials and Methods

### Animals

A total of 24 mice (12 females, 12 males) aged 3-9 months and weighing 20-30g and 7 tree shrews (5 females, 2 males), aged 4 to 33 months and weighing 130-285g were used in this study. **Table 1** reports animals and procedures. This does not include animals that were used for the optimization of these procedures or animals excluded due to lack of reporter expression. The following mouse strains were used in this study: Ai14 (The Jackson Laboratory, Strain #007914 RRID:IMSR_JAX:007914), Chat-Cre (The Jackson Laboratory, Strain #028861; RRID: IMSR_JAX: 028861), Ntsr1-GN209-Cre (Ntsr1-Tg, RRID:MGI:4367044), NPNT-FlpO (The

**Table 1.**
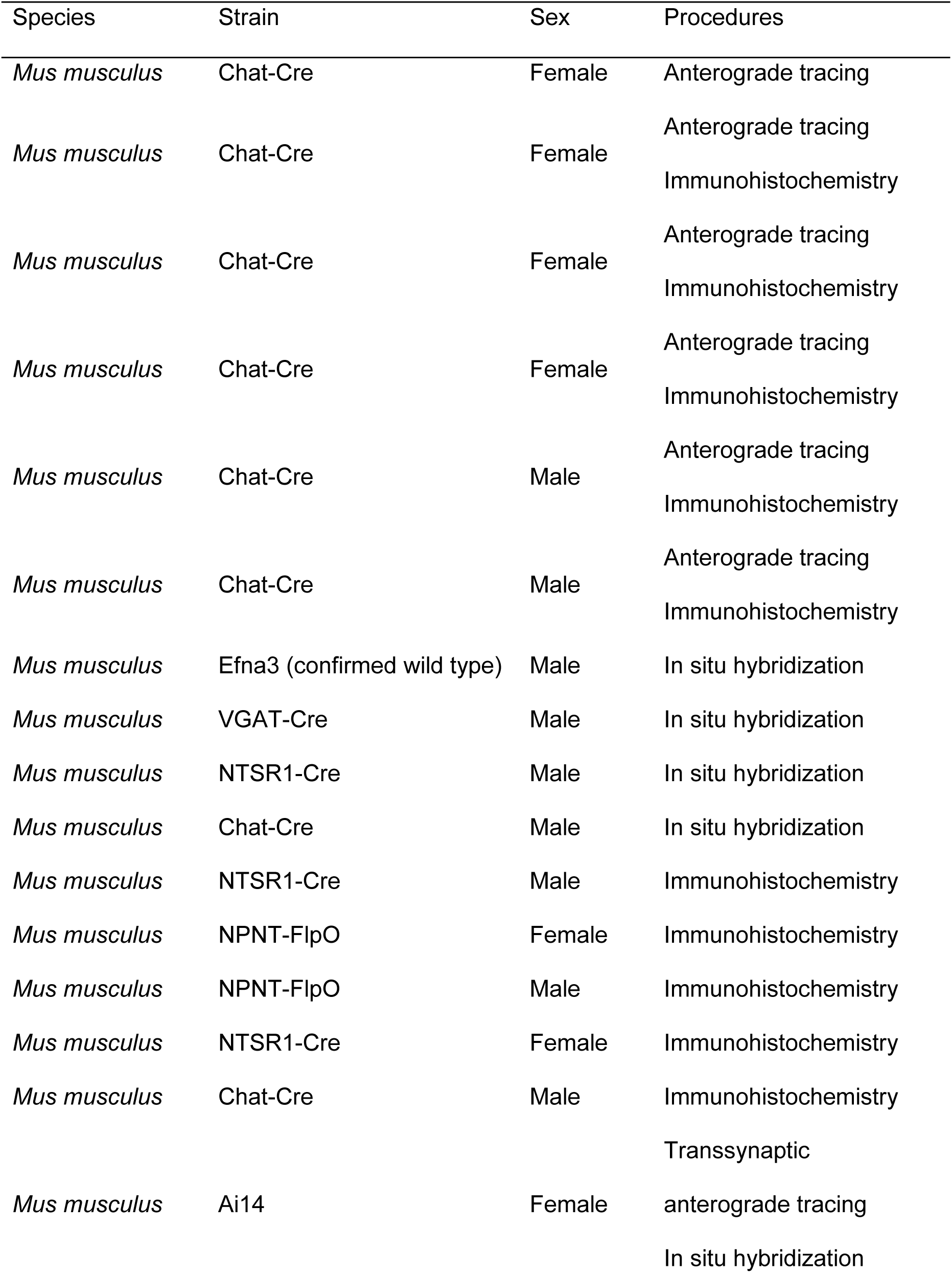
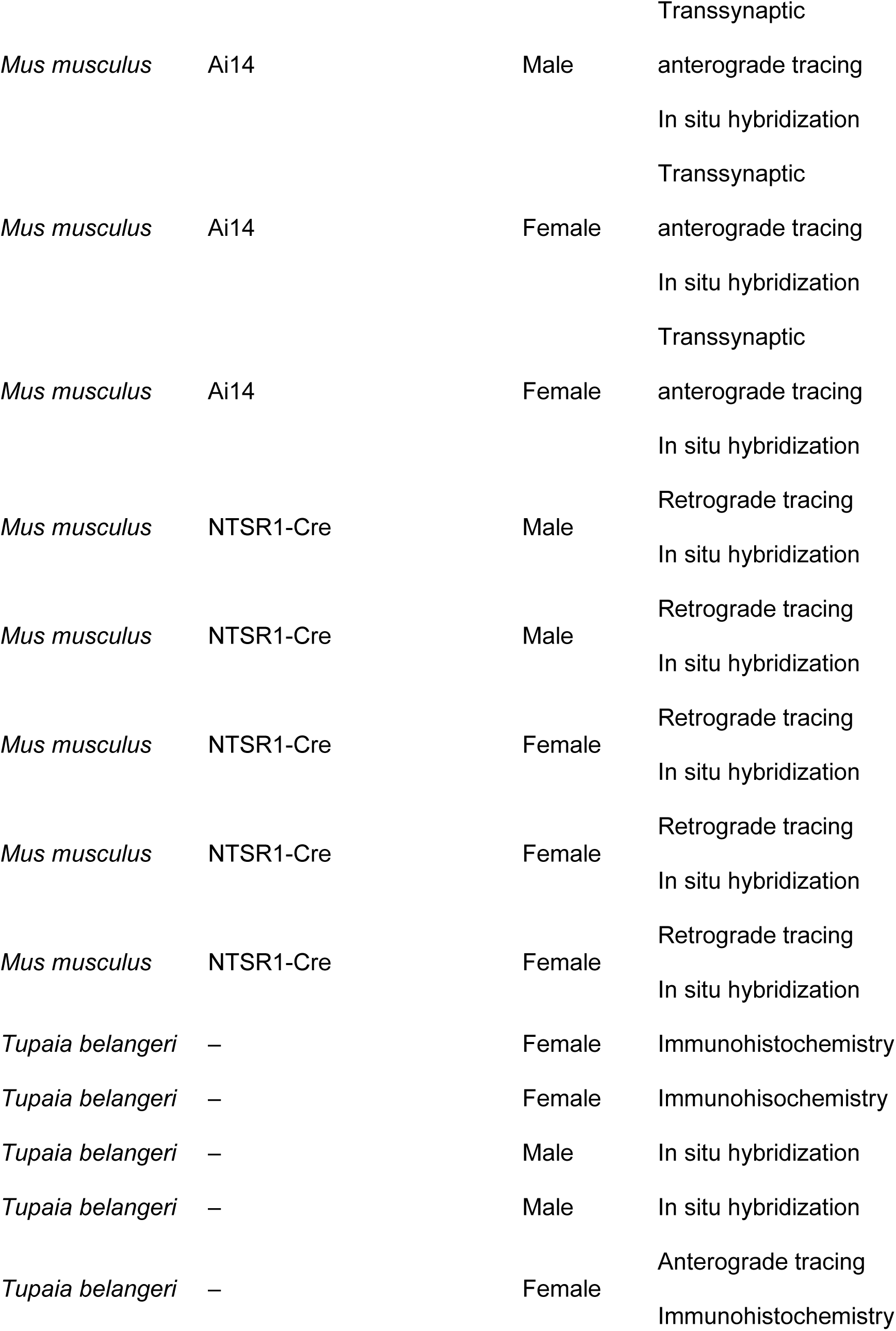
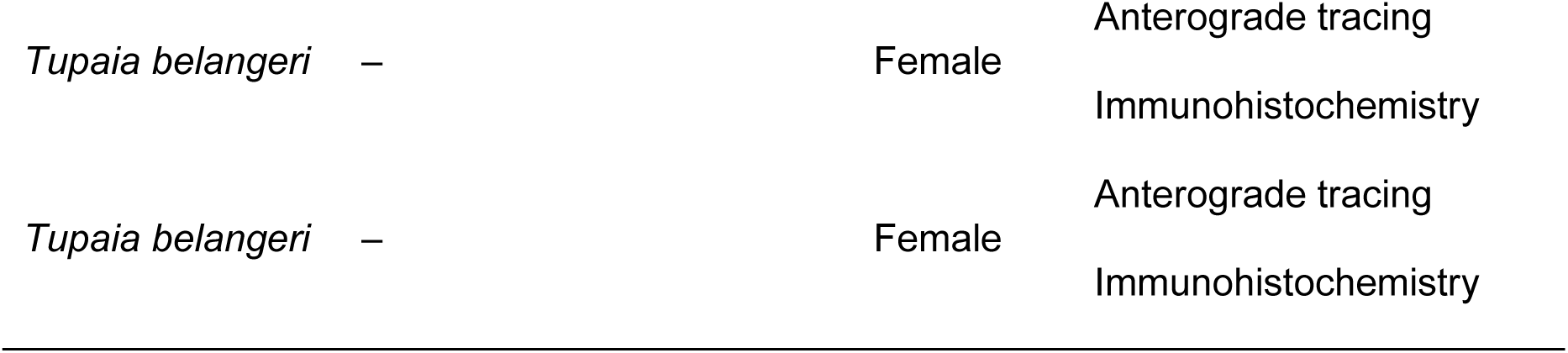
Animals used in this study.

Jackson Laboratory, Strain #034305; RRID:IMSR_JAX:034305), VGAT-Cre (The Jackson Laboratory, Strain #028862; RRID:IMSR_JAX:028862), Isl2-Epha3 (Isl2^tm1(Epha3)Grl^, RRID:MGI:3057124). All mice were kept on a C57BL/6J background. None of the mutations have been reported to produce specific phenotypes other than the expression of Cre in the specified cell-types. Other lines were not carrying the mutant allele. Mice were bred and raised at the University of Michigan and kept on a 12-hour light/dark cycle, with 2-5 animals housed per cage prior to and after surgery. Tree shrews were either provided by the Max Planck Institute or bred and raised at the University of Michigan and kept on a 12-hour light/dark cycle, single housed prior to and after surgery. All experimental procedures were approved by the Institutional Animal Care & Use Committee at University of Michigan.

### Human specimens

De-identified postmortem human brain samples containing the LGN were obtained under institutional policies for exempt, non-human-subjects research. Specimens were provided by the University of Michigan Brain Bank at Michigan Medicine and the NIH NeuroBioBank in Bethesda, Maryland, USA. The study included tissue from 6 donors, consisting of three males (average age: 57.67 ± 5.24) and three females (average age: 56.34 ± 5.58).

### Ethics approval and consent to participate

De-identified postmortem human tissue samples were obtained from established brain bank repositories. These specimens were provided as non-human-subjects research/exempt under institutional policy. Donor consent for tissue collection and research use was obtained by the respective brain bank repositories according to their institutional protocols.

### Stereotaxic injection

Mice were anesthetized with isoflurane (5% induction; 2% maintenance in oxygen at 0.5 L/min) and prepared for surgery. Heads were shaved, a preventive analgesic (Carprofen, 5 mg/kg) was administered, topical lidocaine (2%) at the surgical site, and eye ointment were applied. Animals were then transferred and mounted on a stereotaxic frame, and the surgical site was disinfected using alternating rounds of ethanol and povidone-iodine solution. Temperature was maintained using a feedback temperature controller and a heating pad. Upon confirmation of reflexes loss, an incision was made above the lambda point, and the periosteum was removed. A craniotomy was performed using a microdrill (Foredom Electric, model #K.1070) at the following coordinates: for the SC, 0.5 mm anterior from the lambdoid suture and 0.5mm lateral from the sagittal suture; for the dLGN, 2.2 mm posterior from bregma and 2.25 mm lateral from the sagittal suture. Viral vectors were injected at 1.2 mm (for the SC) and 2.45 mm (for the dLGN) from the brain surface using a nanoliter injector (Drummond Scientific, catalog #3-000-204) with a pulled glass capillary. To achieve the final target volume of 50 nL for the SC and 25 nL for the dLGN, 2.3 nL pulses were delivered every 10 s and the capillary was removed after 10 min post injection to reduce tract labeling. Once the capillary was removed, the craniotomy was closed using bone wax, and the skin was sutured. Incubation time was a minimum of 3 weeks for monosynaptic anterograde and retrograde transport and a minimum of 4 weeks for transsynaptic anterograde transport.

Tree shrews were anesthetized with isoflurane (5% induction; 2% maintenance in oxygen at 0.5 L/min) and prepared for surgery. Heads were shaved, a preventive analgesic (Carprofen, 5 mg/kg), topical lidocaine (2%) at the surgical site, atropine (0.5 mg/kg), and midazolam (2 mg/kg) were administered, and eye ointment was applied. Animals were then transferred and mounted on a stereotaxic frame, and the surgical site was disinfected using alternating rounds of ethanol and povidone-iodine solution. Temperature was maintained using a feedback temperature controller and a heating pad. Temperature, breath rate, heart rate, and mucous membrane color were recorded every 15 min throughout the procedure. Upon confirmation of reflexes loss, an incision was made above the lambda point, and the periosteum was removed. A craniotomy was performed using a microdrill at the following coordinates to target the SC: 0.5mm anterior from the lambdoid suture and 2 mm lateral from the sagittal suture. Viral vectors were injected 3.5 mm ventral to the brain surface using a nanoliter syringe pump (Harvard Biosciences, catalog #788130) with a 5 μL glass syringe (Hamilton Company, product #87908). To achieve the final target volume of 500 nL, viral vectors were delivered with a flow rate of 100 nL/min for 5 min or 250 nL/min for 2 min and the syringe was removed after 5 min post injection to reduce tract labeling. Once the capillary was removed, the craniotomy was closed using bone wax, and the skin was sutured. Incubation time was a minimum of 3 weeks for monosynaptic anterograde transport.

### Perfusion and tissue collection

For experiments that required fixed tissue, animals received a lethal dose of pentobarbital (Euthasol, Virbac) and were transcardially perfused with phosphate buffered saline (PBS, 1x) followed by neutral buffered formalin (NBF, 10%). Brains were harvested and postfixed overnight in 10% NBF at 4 °C. The next day, brains were washed 3 times with 1x PBS, embedded in 5% agarose, and sectioned coronally into 50 μm sections using a vibratome (VT1000, Leica). Tissue sections were stored in well plates containing 1x PBS at 4 °C until used for experiments.

For experiments that required fresh frozen tissue, animals received a lethal dose of pentobarbital (Euthasol, Virbac). Upon the loss of reflexes, brains were harvested, flash frozen on dry ice, and left to completely freeze in-70 °C overnight. The following day, brains were sectioned coronally into 20 μm sections using a cryostat (CM1860, Leica) and mounted directly onto microscope slides (Fisherbrand, Superfrost Plus) and stored at-70 °C until used for experiments.

### Immunohistochemistry (IHC)

Fixed tissue sections were incubated for 1 h at room temperature free floating in a well plate containing 500-1000 μL of blocking buffer solution (normal donkey serum 10%, Triton X-100 0.5%, all in 1x PBS) before incubating in 500-1000 μL of a primary antibody solution for 2 h at room temperature or overnight at 4 °C. Following primary antibody incubation, samples were washed 3 times for 5 min with 1x PBS and incubated in 500-1000 μL of a secondary antibody solution for 1 h and 30 min at room temperature or overnight at 4 °C. All antibodies were diluted using an incubation buffer (normal donkey serum 5%, Triton X-100 0.025%, all in PBS) and their dilutions are reported in **Table 2**. Sections were mounted on microscope slides (Fisherbrand, Superfrost Plus) and counterstained using a mounting medium containing DAPI (SouthernBiotech, DAPI Fluoromount-G).

**Table 2.**
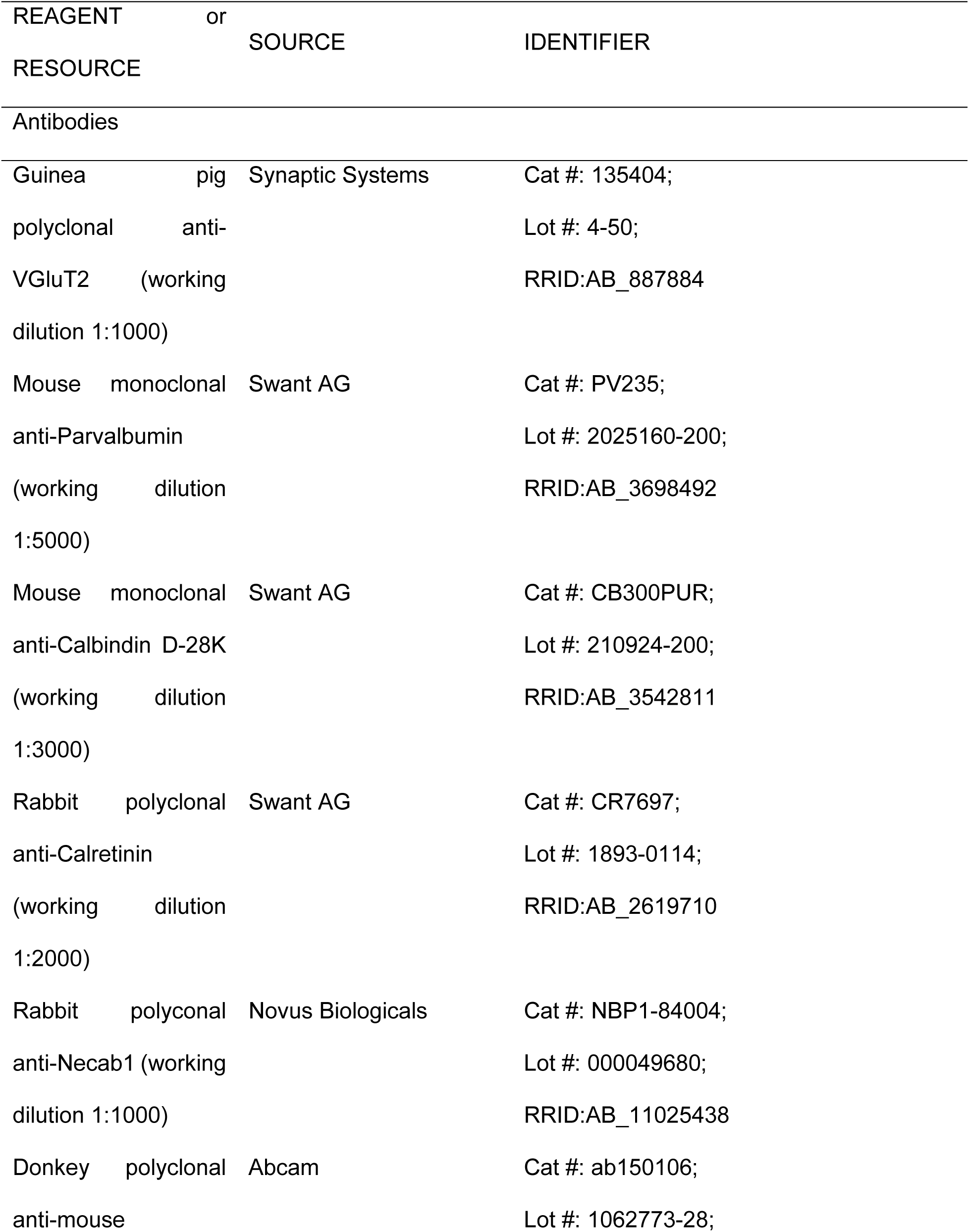
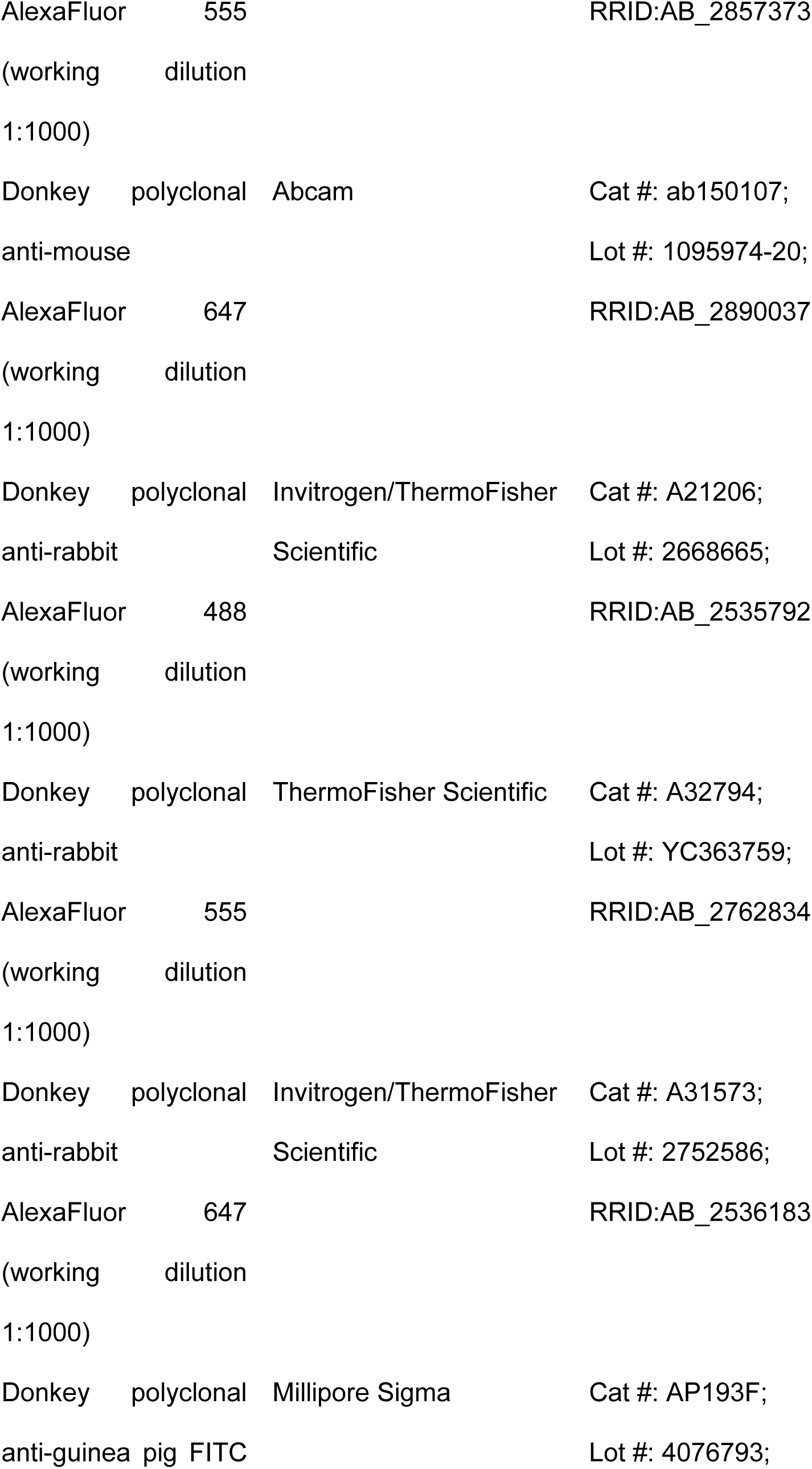
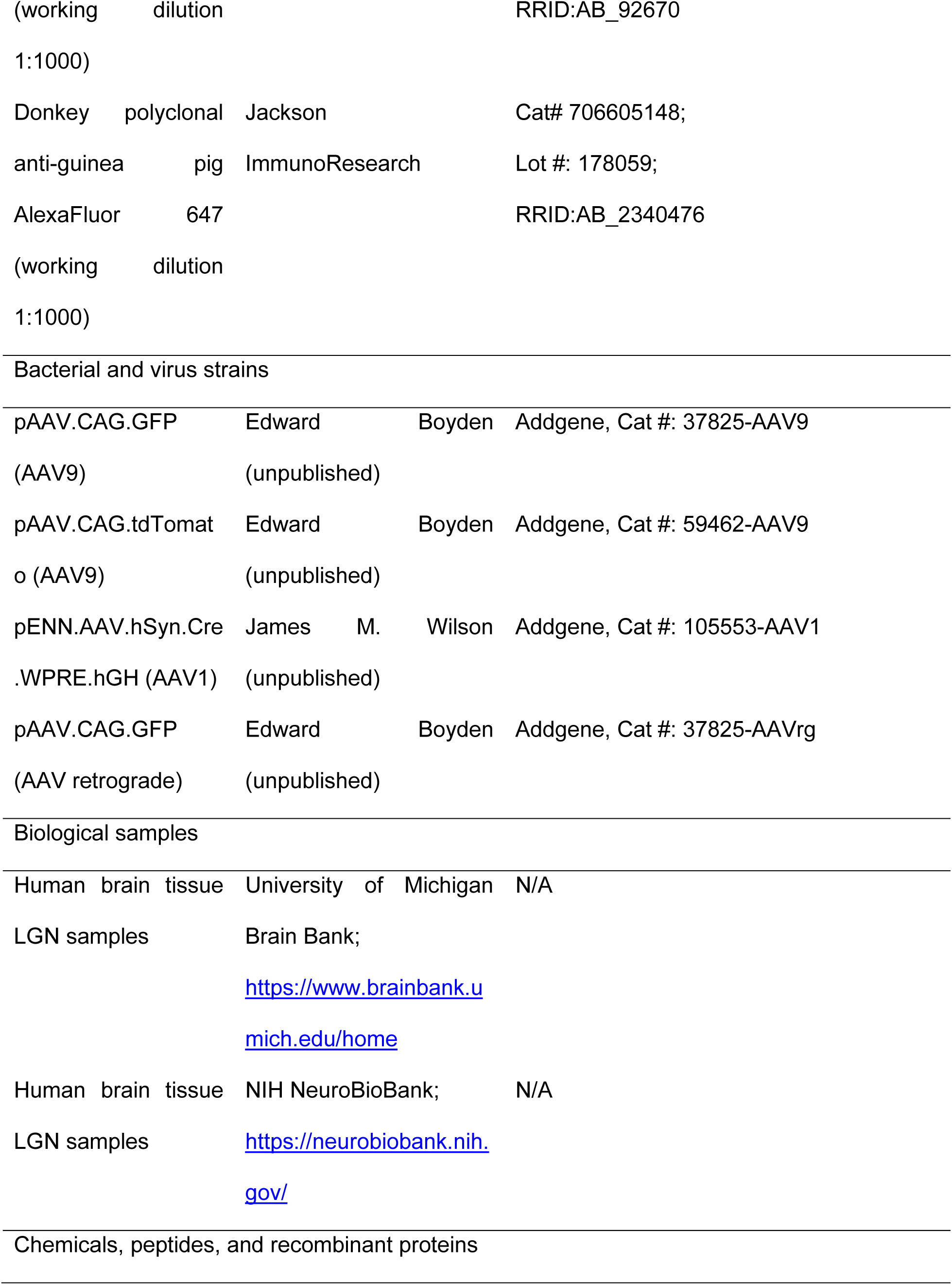
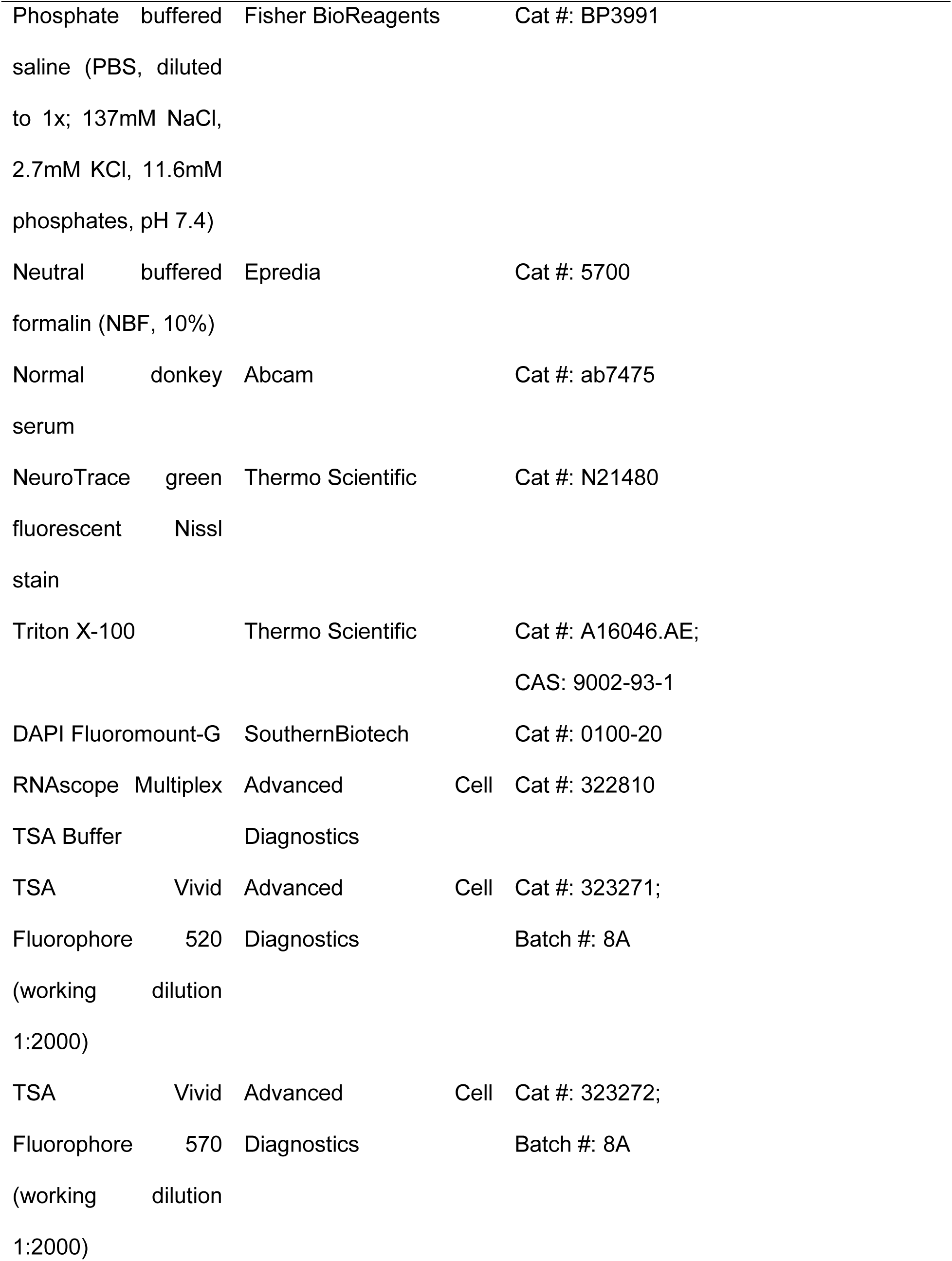
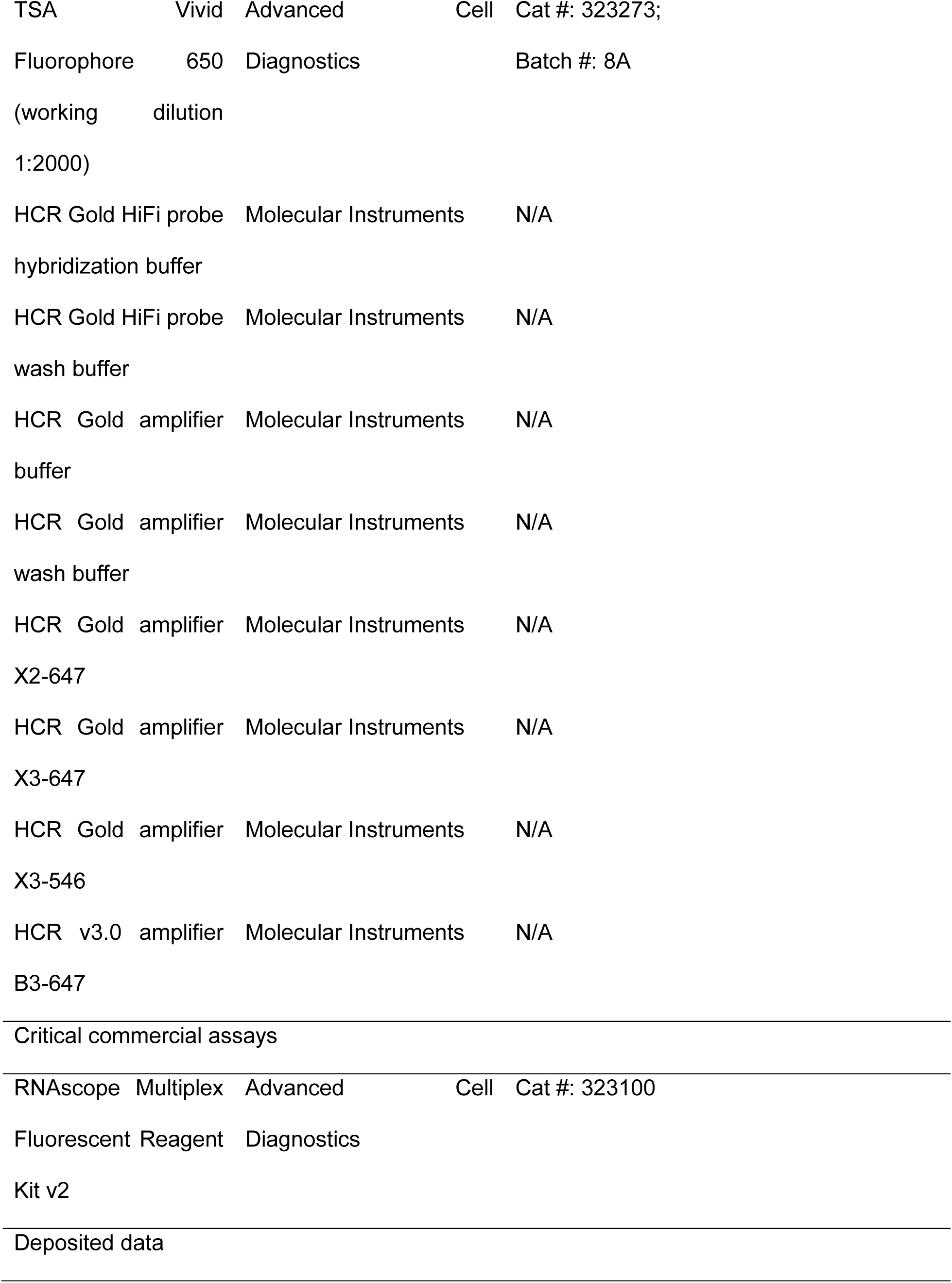
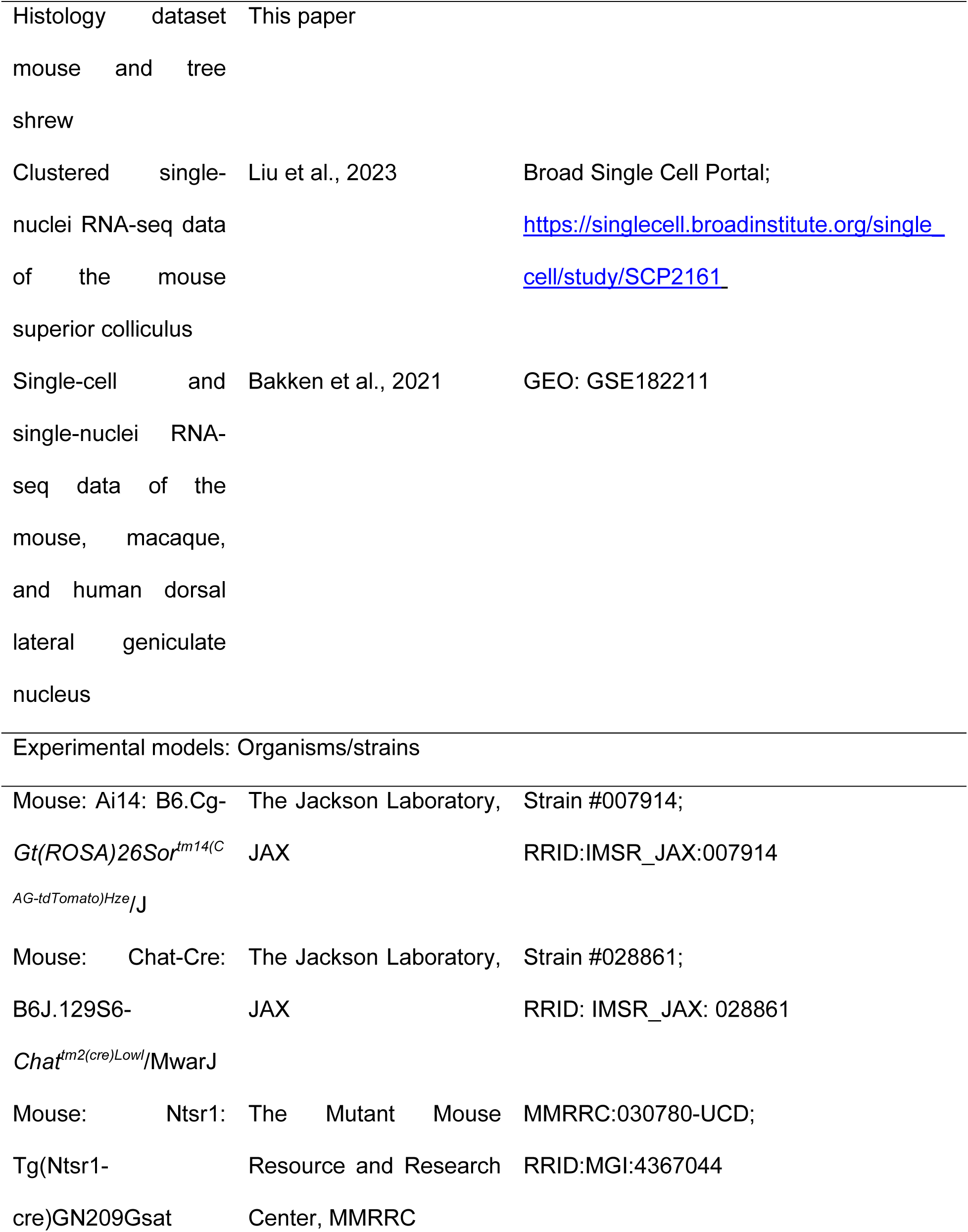
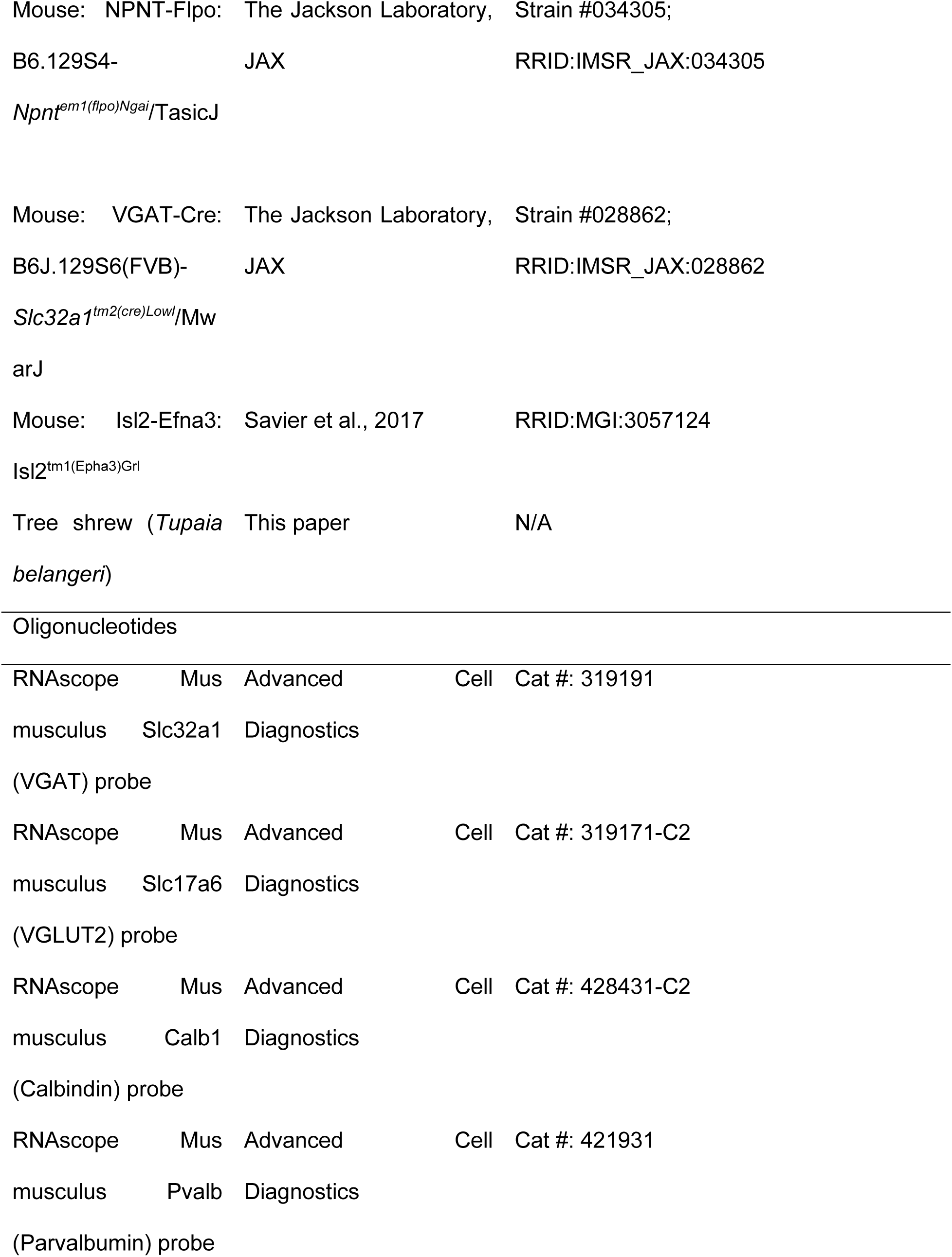
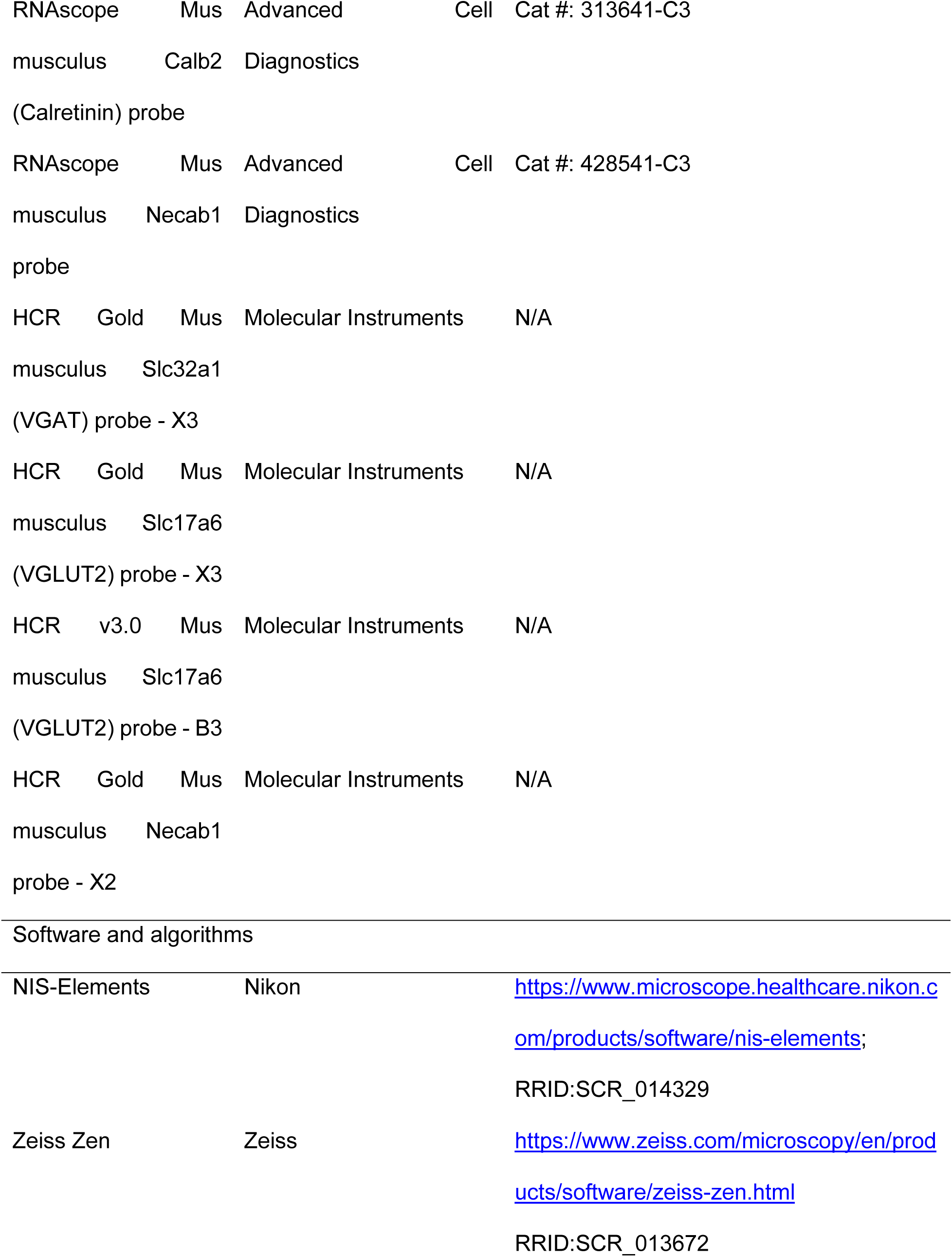
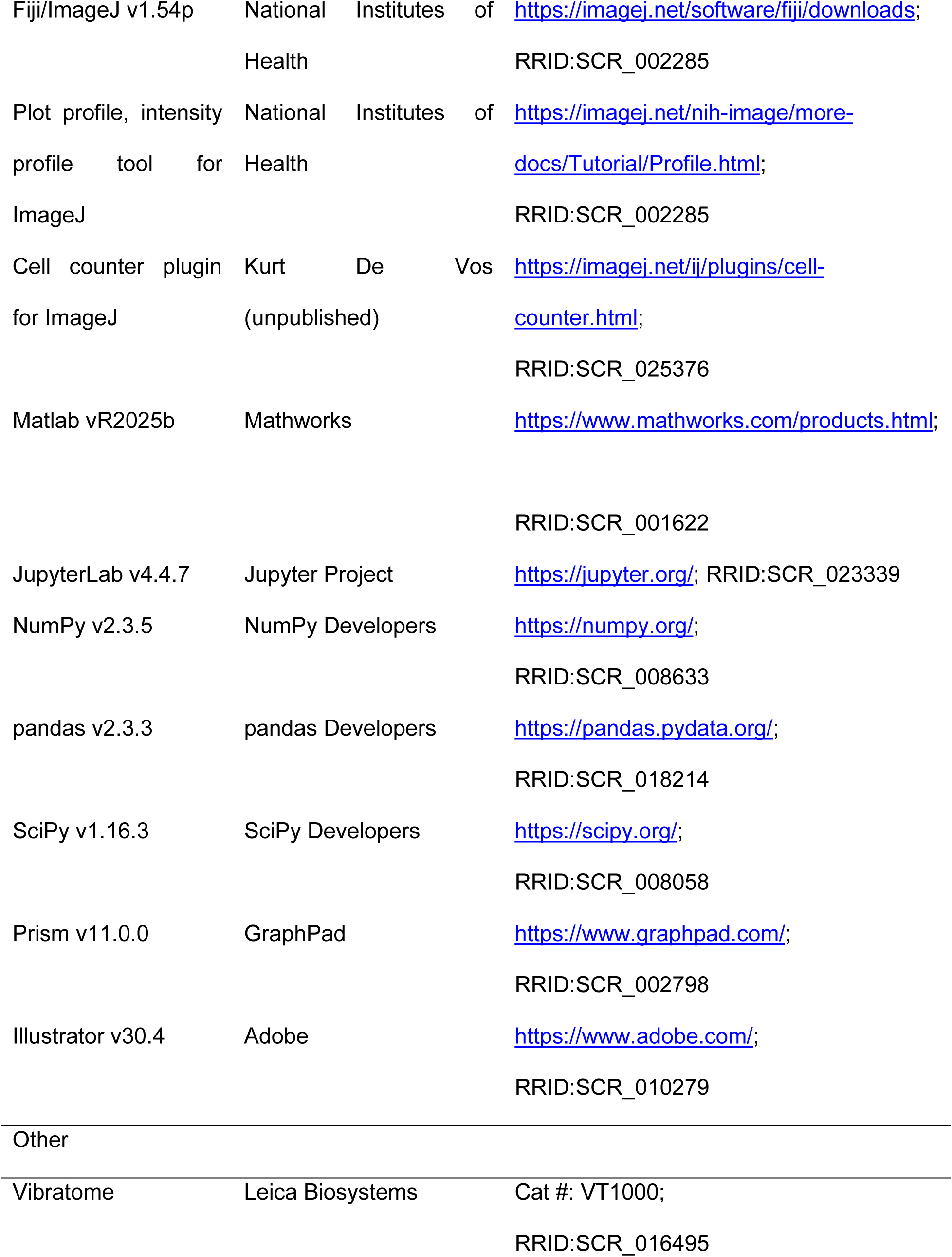
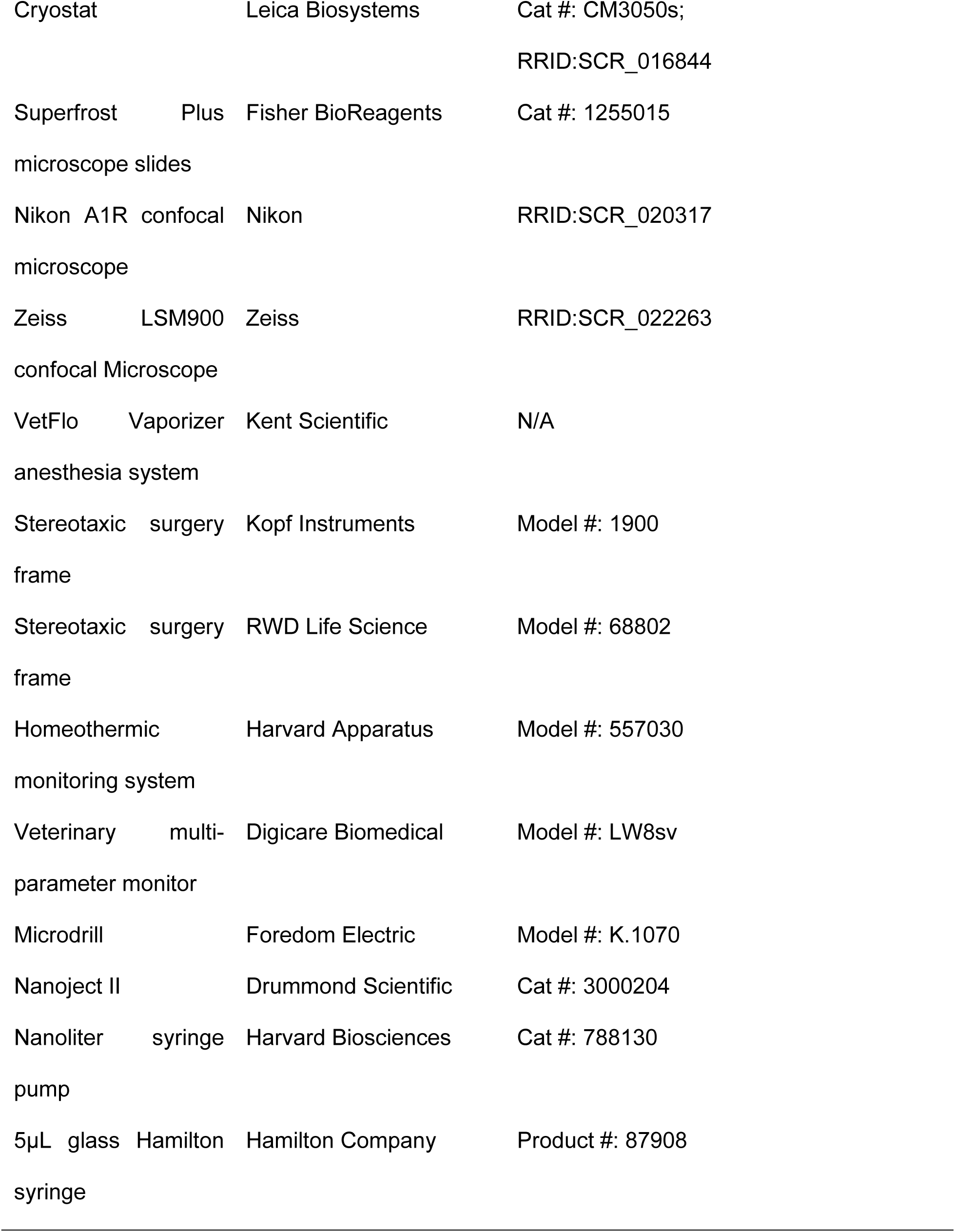
Key resources used in this study.

### Human tissue processing and immunostaining

Postmortem human brain tissue was dissected at autopsy and fixed in formalin. Tissue blocks containing the LGN were selected, embedded in paraffin, and cut into 10 µm thick sections. For immunohistochemical analysis, sections were deparaffinized and rehydrated through a graded ethanol series according to previously established protocols (Werneburg et al., 2017). Antigen retrieval was carried out by heating slides for 10 min in Liberate Antibody Binding Solution (L.A.B. Solution, Polysciences) using a Coplin jar. Following antigen retrieval, sections were blocked and permeabilized for 30 min at room temperature in 10% normal donkey serum, (MilliporeSigma) containing 0.3% Triton X-100 (Sigma-Aldrich) in 0.1 M phosphate buffer. Primary antibody was diluted in the same blocking solution and applied overnight at 4 °C. The primary antibody was rabbit anti-Necab1 (NOVUS, NBP1-84004, 1:1000, RRID: AB_11025438). After primary antibody incubation, sections were washed and incubated for 1 h at room temperature with anti-rabbit-Alexa Fluor–conjugated secondary antibodies (Thermo Fisher Scientific). Sections were then washed and incubated for 20 min (1:300 in 1x PBS) with A488-coupled fluorescent Nissl stain (Thermo Fisher, N21480) according to the manufacturer’s instructions. Following Nissl staining, sections were washed again with 0.1 M phosphate buffer and coverslipped with a mounting medium containing DAPI for nuclear counterstaining (Fluoroshield, MilliporeSigma).

### Fluorescent in situ hybridization (FISH)

Fresh frozen tissue sections were postfixed in 10% NBF for 1 h at 4 °C, followed by dehydration in increasing concentrations of ethanol (50-, 70-, 100%). RNAscope Multiplex V2 (catalog #323100, Advanced Cell Diagnostics) was then performed on sections following the manufacturer’s protease-free protocol. Briefly, sections were treated with 5% hydrogen peroxide to quench endogenous peroxidase activity, then target retrieval was performed using 1x Target Retrieval Reagent (catalog #322000, Advanced Cell Diagnostics) for 10 min at 100 °C, and finally sections were treated with Manual Pretreat Pro (catalog #323100, Advanced Cell Diagnostics) for 30 min at 40 °C. Endogenous mRNA was then hybridized with target probes for 2 h. Following hybridization, probes were amplified with Amps 1-3, and the signal was developed using HRP C1, C2, and C3 followed by binding of the respective fluorophores (TSA Vivid 520/570/650, all diluted 1:2000) and blocked using HRP Blocker between channels. Sections were washed 2 times for 5 min, after each step using a wash buffer (0.1x saline sodium citrate, 0.03% lithium dodecyl sulfate). The probes used in this study are listed in **Table 2**. After the assay, sections were counterstained using a mounting medium containing DAPI (SouthernBiotech, DAPI Fluoromount-G).

### Viral tracing histology

To combine viral tracing with IHC, the aforementioned protocol was followed using 50 μm fixed tissue sections. To combine viral tracing with mRNA-FISH, Molecular Instruments HCR Gold mRNA-FISH was performed on free-floating 50 μm fixed tissue sections using the manufacturer’s instructions. Briefly, samples were dehydrated in increasing concentrations of ethanol (50-, 70-, 100%) and rehydrated in 1x PBS. Next, samples were pre-hybridized using Molecular Instruments HiFi probe hybridization buffer for 30 min at 37 °C. Then, samples were added to probe solutions (10 μL of HCR HiFi probe in 500 μL of HiFi probe hybridization buffer) and allowed to hybridize overnight at 37 °C. After probe hybridization, samples were washed 4 times for 15 min each wash with 1x HiFi probe wash buffer at 37 °C followed by pre-amplification in HCR Gold amplifier buffer for 30 min at room temperature. The probes used in this study are listed in **Table 2**. Amplifier solutions were added to the samples (10 μL of HCR Gold amplifier hairpin 1 and 10 μL of HCR Gold amplifier hairpin 2 in 500 μL of HCR Gold amplifier buffer) and allowed to incubate overnight at room temperature on an orbital shaker. Samples were washed 4 times for 15 min each wash with 1x HCR Gold amplifier wash buffer at room temperature, mounted on microscope slides (Fisherbrand, Superfrost Plus), and counterstained using a mounting medium containing DAPI (SouthernBiotech, DAPI Fluoromount-G).

### Image acquisition

Samples were imaged using a Nikon A1R confocal microscope equipped with 10x/0.45 NA Plan Apo λ and 20x/0.75 NA Plan Apo λ objectives. One representative image of the human dLGN was collected using a Zeiss LSM900 confocal microscope equipped with 20x/0.8 M27 Plan-Apochromat with Zen-Blue software. All other images were acquired using Nikon NIS-Elements software as either Z-stacks, tiled images, or tiled Z-stacks. 10x Z-stacks were acquired using a 3.2- to 3.75 μm step size and 20x Z-stacks were acquired using a 0.95- to 1.1 μm step size. All images were acquired using galvo scanning at 1024 × 1024 pixel resolution.

## Quantification and statistical analysis

### Intensity profile quantification

To determine the spatial expression of molecular markers in the dLGN, 10x Z-stacks of the tissue sections were opened in Fiji/ImageJ (RRID:SCR_002285, Version 1.54p, National Institutes of Health), and Z-projections were generated using the maximum intensity. Images were rotated so that the lateral-most edge of the dLGN was oriented perpendicular to the lower image boundary. **Figure S1** illustrates this protocol in greater detail. A rectangular region of interest measuring 100 μm in height and spanning the full width of the dLGN was then drawn across the region. Fluorescence intensity profiles were extracted from this region using the Plot Profile tool in Fiji/ImageJ.

For comparison across sections and experimental groups, the distance of each profile was normalized using min-max normalization, where 0 corresponds to the medial-most edge and 1 corresponds to the lateral-most edge. Intensity profiles were then interpolated to 1000 equally spaced points using the interp1d function in SciPy (RRID:SCR_008058). Interpolated profiles were used to calculate mean intensity profiles for each dataset and to compare spatial expression patterns across groups.

To bin the intensity by the region of the mouse dLGN, a curve was fitted to the averaged intensity profiles of VGluT2 IHC, which was then used to calculate the maximum of the second derivative to determine the position where the intensity profile most rapidly changed. This point was used as the boundary between the shell and the core of the mouse dLGN (dashed grey line, Fig. 1A, bottom right). For the tree shrew, we used a similar method by first gathering the intensity profiles of VGluT2 IHC, averaging these (Fig. 1B, bottom right), then fitting a curve to the resulting data, and finding the extrema of the first derivative to account for each layer and ILZ (dashed grey lines, Fig. 1B, bottom right). Using these boundaries, we binned and subsequently averaged the data from each intensity profile.

**Figure 1.**
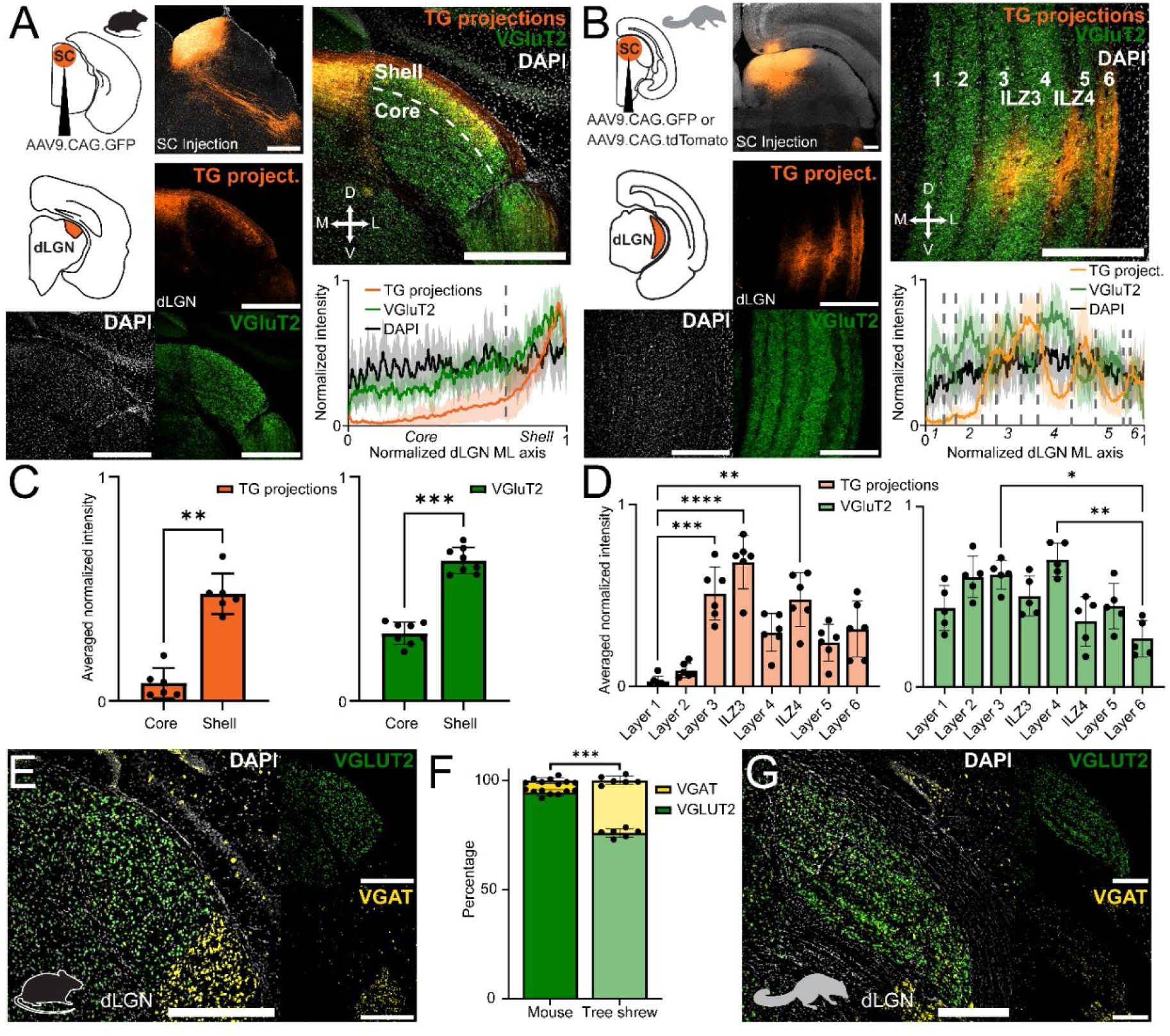
Anatomical tracing of the tectogeniculate projections and quantification of neurotransmitter distribution in the mouse and tree shrew dLGN. A. Anterograde tracing in the mouse dLGN using an injection of AAV9.CAG.GFP in the SC. IHC of VGluT2 in the mouse dLGN (green), fibers from the SC (orange), and DAPI counterstaining (white). Normalized mean intensity profile with shaded standard deviation of a 100 µm rectangular ROI spanning across the medial-lateral axis of the dLGN (bottom right; n = 6 animals, 6 sections for tectogeniculate projections data; n = 5 animals, 8 sections for VGluT2 data). B. Same as A but for the tree shrew (n = 3 animals, 6 sections for tectogeniculate projections data; n = 5 animals, 5 sections for VGluT2 data). C. Averaged normalized intensity binned for “core” and “shell” of the mouse dLGN using VGluT2-defined regions (Mann-Whitney test; p ≤ 0.01 (**) and ≤ 0.001 (***); tectogeniculate projections between core and shell, p = 0.002; VGluT2 intensity between core and shell, p = 0.0002; n = 6 animals, 6 sections for tectogeniculate projections data; n = 5 animals, 8 sections for VGluT2 data). D. Averaged normalized intensity binned by layers and ILZs using VGluT2-defined boundaries (Kruskal-Wallis test with Dunn’s test for multiple comparison corrections; p ≤ 0.05 (*), ≤ 0.01 (**), ≤ 0.001 (***), and ≤ 0.0001 (****); Tectogeniculate projections data comparing the mean rank of each column to the mean rank of Layer 1 using the Kruskal-Wallis test, p < 0.0001; Dunn’s test comparing between layers 1 and 3, p = 0.0005; between layers 1 and ILZ3, p < 0.0001; between layers 1 and ILZ4, p = 0.0014; between layers 1 and 6, p = 0.0618; VGluT2 data comparing the mean rank of each column to the mean rank of every other column Kruskal-Wallis test, p = 0.0007; Dunn’s test comparing between layers 3 and 6, p = 0.0202; between layers 4 and 6, p = 0.0020; n = 3 animals, 6 sections for tectogeniculate projections data; n = 5 animals, 5 sections for VGluT2 data). E and G. RNAscope FISH staining of a mouse (E) and tree shrew (G) dLGN for VGLUT2 (green) and VGAT (yellow). F. Bar graph to compare the ratio of VGLUT2+ to VGAT+ cells in the mouse and the tree shrew using normalized cell count (Mann-Whitney test; p ≤ 0.001 (***); VGLUT+ to VGAT+ between mouse and tree shrew, p = 0.0007; n = 4 animals, 8 sections, 3540 total cells counted, mean ± SD: 442.5 ± 47.64 cells per section for mouse data; n = 2 animals, 6 sections, 5725 total cells counted, mean ± SD: 954.17 ± 67.38 cells per section for tree shrew data). All data presented as mean ± SD. All scale bars are 500 µm.

### Molecular identity quantification for transsynaptic and retrograde labeling

To determine the molecular identity of cells in viral tracing experiments, 20x Z-stacks were opened in Fiji/ImageJ and Z-projections were generated using the maximum intensity. Using the Cell Counter plugin in Fiji/ImageJ, the number of reporter-labeled cells, marker-positive cells, and colocalized cells were manually counted. Reporter-positive cells were identified by the fluorescent reporter signal that filled the soma. Cells were classified as marker-positive if puncta formed a ring-like pattern corresponding to the outline of a cell body. Cells were considered co-expressing if they met both criteria. To determine the ratio of excitation to inhibition, the number of VGLUT2+ and VGAT+ cells were counted in each section and then the sum of the two was determined to be the total number of cells. Then the number of VGLUT2+ cells was divided by the total number of cells to determine the ratio.

### Experimental design and statistical analysis

Statistical tests used for each analysis are indicated in the corresponding figure captions, along with the number of animals, tissue sections, and cells analyzed. A *p*-value ≤ 0.05 was considered statistically significant. Due to the sample size, normality could not be determined, and all statistical tests were nonparametric. No statistical methods were used to predetermine sample size, but the sample size is in accordance with other similar studies. Due to the sample size, data collected from both female and male animals were pooled together, but no sex-related differences were noted. When comparing multiple markers or conditions, sections missing one of the compared conditions were excluded only for that specific comparison.

## Results

### The mouse and tree shrew dLGN exhibit distinct organizations of tectorecipient subdivisions and neurotransmitter distribution

To assess the organization of tectorecipient subdivisions within the dLGN, an anterograde viral tracer was injected in the SC of both the mouse and the tree shrew. Since VGluT2 protein is found in the layers of the tree shrew dLGN but not the ILZs due to lack of inputs from the retina (Balaram et al., 2015), we developed a method of determining subdivisions within the mouse and tree shrew dLGN utilizing the intensity profiles of VGluT2 IHC staining (Fig. 1A bottom right, for details see Methods and **Figure S1)**. In the mouse, tectogeniculate projections are restricted to the shell of the dLGN (Fig. 1C, left; mean ± SD: core, 0.08 ± 0.07; shell, 0.48 ± 0.09) and the shell has a higher expression of VGluT2 expression than the core (Fig. 1C, right; mean ± SD: core, 0.30 ± 0.05; shell, 0.63 ± 0.06). In the tree shrew, tectogeniculate projections were restricted to layers 3 and 6, as well as the ILZs between layers 3 and 4 (ILZ3) and layers 4 and 5 (ILZ4) with the greatest intensity found within ILZ3 (Fig. 1D, left; mean ± SD: layer 1, 0.03 ± 0.03; layer 3, 0.51 ± 0.15; ILZ3, 0.68 ± 0.15; ILZ4, 0.48 ± 0.15; layer 6, 0.32 ± 0.15). Intensity of tectogeniculate projections depended on the location of the injection site as well as the AP axis of the dLGN (see **Figure S2**). VGluT2 protein is distributed relatively evenly across the dLGN of the tree shrew, with lower expression found in the ILZs as well as layer 6 (Balaram et al., 2015) (Fig. 1D, right mean ± SD: ILZ3, 0.50 ± 0.11; ILZ4, 0.36 ± 0.14; layer 6, 0.27 ± 0.10). In both the mouse and the tree shrew, there were no contralateral projections observed (data not shown). Overall, our results show that the mouse and the tree shrew dLGN exhibit discrete organization of tectogeniculate projections, with restriction to the shell in the mouse and the targeting of distinct laminae in the tree shrew. To address other core organization principles, we next examined the ratio of excitatory to inhibitory neurons in the dLGN of the mouse and the tree shrew using RNAscope FISH for VGLUT2 and VGAT mRNA. Previous studies have shown little mRNA expression for VGLUT1 in the tree shrew (Balaram et al., 2015) and VGLUT3 in the mouse (Fremeau et al., 2004) dLGN, so we focused on VGLUT2 to determine the proportion of GABAergic to glutamatergic neurons. This resulted in the finding that the tree shrew shows lower levels of VGLUT2+ cells or higher levels of relative inhibition. In the mouse, a sharp transition can be observed between the dorsal and the ventral LGN regarding the proportion of GABAergic neurons (Fig 1E). While no patterns in the spatial distribution of GABAergic neurons were observed within the dLGN of both species (Fig. 1E & G), a significant increase in the proportion of GABAergic neurons was observed in tree shrews (GABAergic neuron % mean ± SD: mice, 5.59 ± 1.36%; tree shrews, 24.14 ± 1.92%; Fig. 1F). To further investigate potential subdivisions and regional specialization of the mouse and tree shrew dLGN, we next addressed the distribution and conservation of previously used molecular markers.

### Classical markers reveal species-specific molecular organization in the mouse and tree shrew dLGN

To determine the distribution and conservation of previously identified molecular markers in the mouse and tree shrew dLGN, we performed both IHC and RNAscope FISH of calcium binding proteins parvalbumin (PV, Pvalb), calbindin (CB, Calb1), and calretinin (CR, Calb2). Our findings indicate that these markers show a variable conservation between the mouse and tree shrew dLGN. In the mouse dLGN, PV showed similar expression throughout the shell and the core (Fig. 2A, red; mean ± SD: core, 0.58 ± 0.08; shell, 0.68 ± 0.09). CB protein has been found in the shell of the mouse dLGN (Grubb and Thompson, 2004) and our results agree with this finding (Fig. 2A, magenta; mean ± SD: core, 0.26 ± 0.05; shell, 0.58 ± 0.06). A similar expression pattern was found for CR in the shell of the mouse dLGN (Fig. 2A, blue; mean ± SD: core, 0.28 ± 0.02; shell, 0.59 ± 0.09). However, CR expression cannot be found in somas within the dLGN, while it can be observed in the lateral posterior nucleus (LP), intergeniculate leaflet (IGL), and ventral lateral geniculate nucleus (vLGN). In the tree shrew, PV expression is found throughout all layers with the lowest expression in layer 6 and ILZ4 (the regions with tectogeniculate projections) and highest in layers 2 and 4 (Fig. 2B, red; mean ± SD: layer 2, 0.55 ± 0.09; layer 4, 0.65 ± 0.09; ILZ4, 0.33 ± 0.07; layer 6, 0.25 ± 0.05), contrasting with findings in the mouse. CB is found to have the lowest expression in layers 1 and 2 with a sharp increase in expression in layer 3 that slowly decreases as the layers progress along the mediolateral axis (Fig. 2B, magenta; mean ± SD: layer 1, 0.23 ± 0.07; layer 2, 0.23 ± 0.03; layer 3, 0.56 ± 0.15; layer 6, 0.38 ± 0.18). CR is also found to have the highest expression in layer 3 and the lowest expression in layer 6; however, there is expression present within each of the layers (Fig. 2B, blue; mean ± SD: layer 3, 0.67 ± 0.13; layer 6, 0.25 ± 0.06). Differing from the results in the mouse, CR immunoreactive somas could be found in the dLGN of the tree shrew. Examining FISH data within the mouse, Pvalb mRNA shows a similar expression as seen in **Fig. 2A**, however with a slight bias for the core, as seen in the representative photomicrograph (Fig. 2C, red; mean ± SD: core, 0.31 ± 0.06; shell, 0.30 ± 0.03). This is a similar finding to what has been established in the literature, with Pvalb mRNA showing a gradient of expression between the shell and core regions (Bakken et al., 2021). When looking at the intensity profile, however, there seems to be no difference in expression between the shell and the core, possibly due to the sparsely positive cells and autofluorescence of the tissue. Calb1 mRNA shows a bias for the shell of the dLGN in both the photomicrograph and intensity profile (Fig. 2C, magenta; mean ± SD: core, 0.32 ± 0.08; shell, 0.39 ± 0.05), but this bias is not significant. Interestingly, there is no expression of Calb2 (CR) mRNA in the mouse dLGN (Fig. 2C, blue; mean ± SD: core, 0.19 ± 0.06; shell, 0.25 ± 0.08), which is especially noticeable when comparing it to the expression pattern seen in the LP and IGL, and any intensity measured would be attributed to tissue autofluorescence. The tree shrew dLGN shows similar patterns of PVALB mRNA expression as seen in IHC data with high expression in layers 2 and 4 and low expression in layer 6 (Fig. 2D, red; mean ± SD: layer 2, 0.54 ± 0.08; layer 4, 0.51 ± 0.07; layer 6, 0.22 ± 0.06). The same consistency between IHC and FISH data is also seen with CALB1 (CB) with an increase in expression in layer 3 (Fig. 2D, magenta; mean ± SD: layer 1, 0.16 ± 0.11; layer 3, 0.51 ± 0.10; layer 6, 0.16 ± 0.05). In stark contrast to the mouse, CALB2 (CR) mRNA is present in every layer of the tree shrew dLGN with the highest expression in layer 4 and lowest expression in layer 6, much akin to CR immunoreactivity (Fig. 2D, blue; mean ± SD: layer 4, 0.50 ± 0.05; layer 6, 0.18 ± 0.12). Overall, using IHC and mRNA-FISH, we found variable conservation of classical markers between the mouse and tree shrew dLGN, and the distribution of previously identified molecular markers did not overlap neatly with the distribution of tectorecipient inputs in either species. As a result, we then turned to transcriptomic-based approaches to identify conserved molecular markers that are specific to the tectorecipient layers of the dLGN.

**Figure 2.**
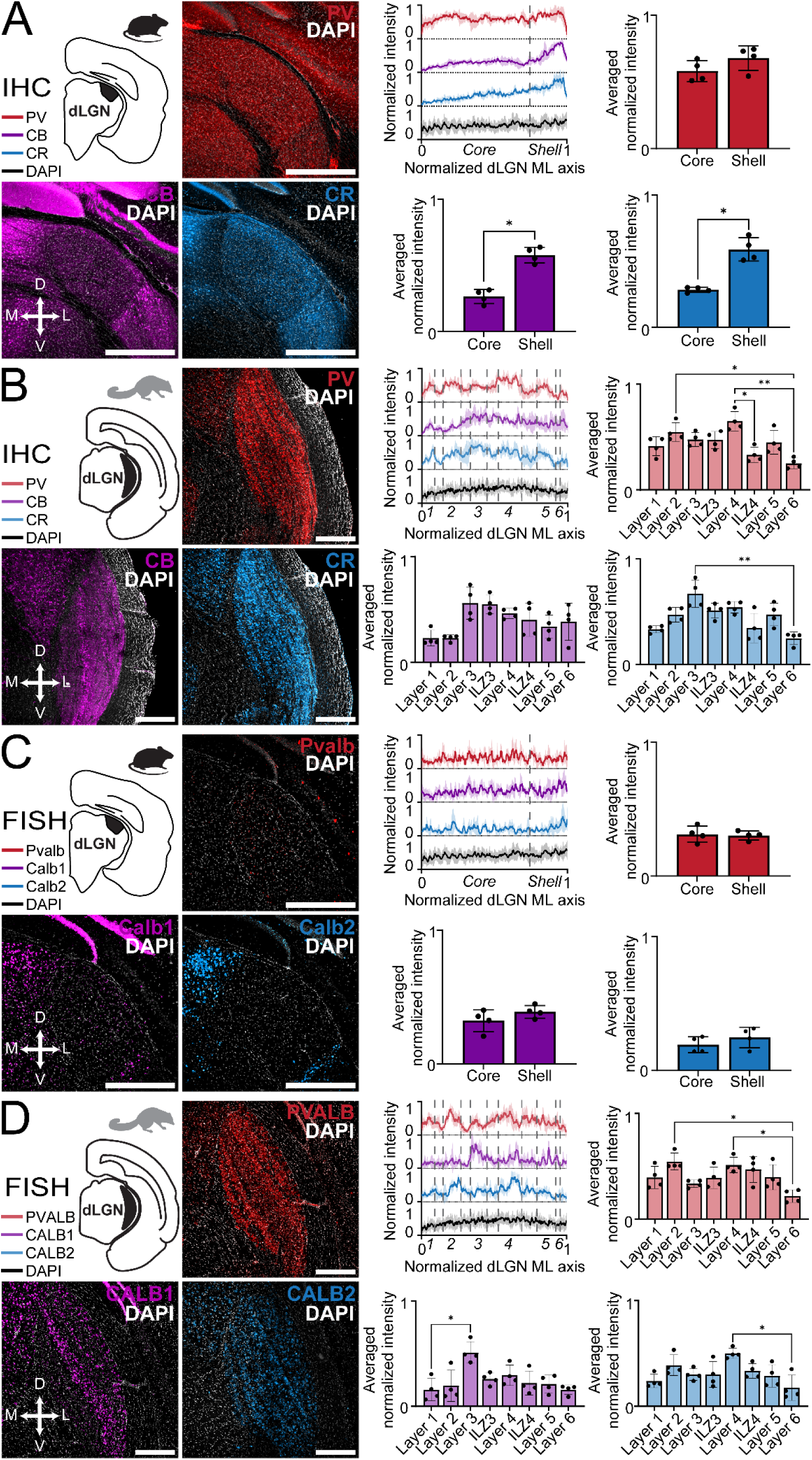
Immunostaining and fluorescent in situ hybridization of calcium binding proteins in the mouse and tree shrew dLGN.A. IHC in the mouse dLGN of parvalbumin (PV, red), calbindin (CB, magenta), and calretinin (CR, blue). Normalized mean intensity profiles with shaded standard deviation of a 100 µm rectangular ROI spanning across the medial-lateral axis of the dLGN. Averaged normalized intensity binned for “core” and “shell” of the mouse dLGN using VGluT2-defined regions from Figure 1 (Mann-Whitney test; p ≤ 0.05 (*); core versus shell: PV, p = 0.1143; CB, p = 0.0286; CR, p = 0.0286; n = 4 animals, 4 sections). **B.** IHC in the tree shrew dLGN of PV (red), CB (magenta), and CR (blue). Normalized mean intensity profiles with shaded standard deviation of a 100 µm rectangular ROI spanning across the medial-lateral axis of the dLGN. Averaged normalized intensity binned for layers and ILZs of the tree shrew dLGN using VGluT2-defined regions from Figure 1 (Kruskal-Wallis test with Dunn’s test for multiple comparison corrections comparing the mean rank of each column with the mean rank of every other column; p ≤ 0.05 (*) and ≤ 0.01 (**); layer comparisons for PV: Kruskal-Wallis test, p = 0.0027; Dunn’s test comparing between layers 2 and 6, p = 0.0492; between layer 4 and ILZ4, p = 0.0380; between layers 4 and layers 6, p = 0.0025; layer comparisons for CB: Kruskal-Wallis test, p = 0.0122; layer comparisons for CR: Kruskal-Wallis test, p = 0.0037; Dunn’s test comparing between layers 3 and 6, p = 0.0062; n = 4 animals, 4 sections). **C.** Same as **A** except for RNAscope FISH (Mann-Whitney test, core versus shell: Pvalb, p > 0.9999; Calb1, p > 0.9999; Calb2, p = 0.3429; n = 4 animals, 4 sections). **D.** Same as **B** except for FISH (layer comparisons for PVALB: Kruskal-Wallis test, p = 0.0155; Dunn’s test comparing between layers 2 and 6, p = 0.0111; between layers 4 and 6, p = 0.0492; layer comparisons for CALB1: Kruskal-Wallis test, p = 0.0237; Dunn’s test comparing between layers 1 and 3, p = 0.0492; layer comparisons for CALB2: Kruskal-Wallis test, p = 0.0236; Dunn’s test comparing between layers 4 and 6, p = 0.0169; n = 2 animals, 4 sections). All data presented as mean ± SD. All scale bars are 500 µm. Necab1 serves as a conserved molecular marker for regions that receive tectogeniculate projections and is present in the dLGN of rodents, tree shrews, and humans.

Given the variable conservation of classical markers and their limited specificity for dLGN regions receiving tectogeniculate projections, we turned to recent RNAseq data to identify a more selective molecular marker. Previous studies found that Necab1 shows a preference for the shell of the mouse dLGN and the K layers of the primate dLGN (Bakken et al., 2021). Leveraging this, we performed IHC to assess the immunoreactivity of Necab1 in the mouse (Fig. 3A), tree shrew (Fig. 3B), and human dLGN (Fig. 3C). Intensity profiles of these IHC experiments revealed discrete regions in the mouse (Fig. 3D) and tree shrew dLGN (Fig. 3E), and specificity for the human dLGN (Fig. 3C and 3F). Photomicrographs from additional human donors can be found in **Figure S3** (n = 6 donors, 6 sections). Necab1 immunoreactivity is restricted to the shell of the mouse dLGN (Fig. S4A, Mann-Whitney test, *p* = 0.0002) and in the tree shrew it is present in layers 3 and 6 as well as ILZ3 and ILZ4 (Fig. S4B, Kruskal-Wallis test, *p* < 0.0001; Dunn’s test comparing between layers 1 and 3, *p* < 0.0001; between layer 1 and ILZ3, *p* = 0.0004; between layer 1 and ILZ4, *p* = 0.0261; between layers 1 and 6, *p* = 0.0414). To confirm the findings from IHC data, we performed RNAscope FISH on mouse dLGN (Fig. 3G, left) and tree shrew dLGN (Fig. 3H, left) sections. FISH of Necab1 in the mouse shows a specificity for the shell region (Fig. 3G). FISH of NECAB1 in the tree shrew shows specificity for layers 3 and 6 and ILZ3 and ILZ4 (Fig. 3H). Following an injection of an anterograde viral tracer in the SC (**Figure 1**), we performed IHC on sections of the dLGN of the mouse (Fig. 3I) and tree shrew (Fig. 3J) to determine if Necab1 expression and tectogeniculate projections overlap. Intensity profiles of both tectogeniculate projections and Necab1 IHC staining in the mouse dLGN show a similar pattern (Fig. 3K, left) and show no statistical difference in intensity (Fig. 3K, right). The same relationship is found in the tree shrew dLGN intensity profiles (Fig. 3L, left) and in the comparison between intensities with a slight difference in layer 3 (Fig. 3L, right). The slight difference in intensity within layer 3 could be attributed to the small bias for the SC to project to ILZ3. Overall, our results show that Necab1 expression, both at the mRNA and protein level, overlaps with tectogeniculate projections in the mouse and tree shrew dLGN. We next compared our new findings with CB expression, a previously identified marker for tectorecipient layers of the dLGN.

**Figure 3.**
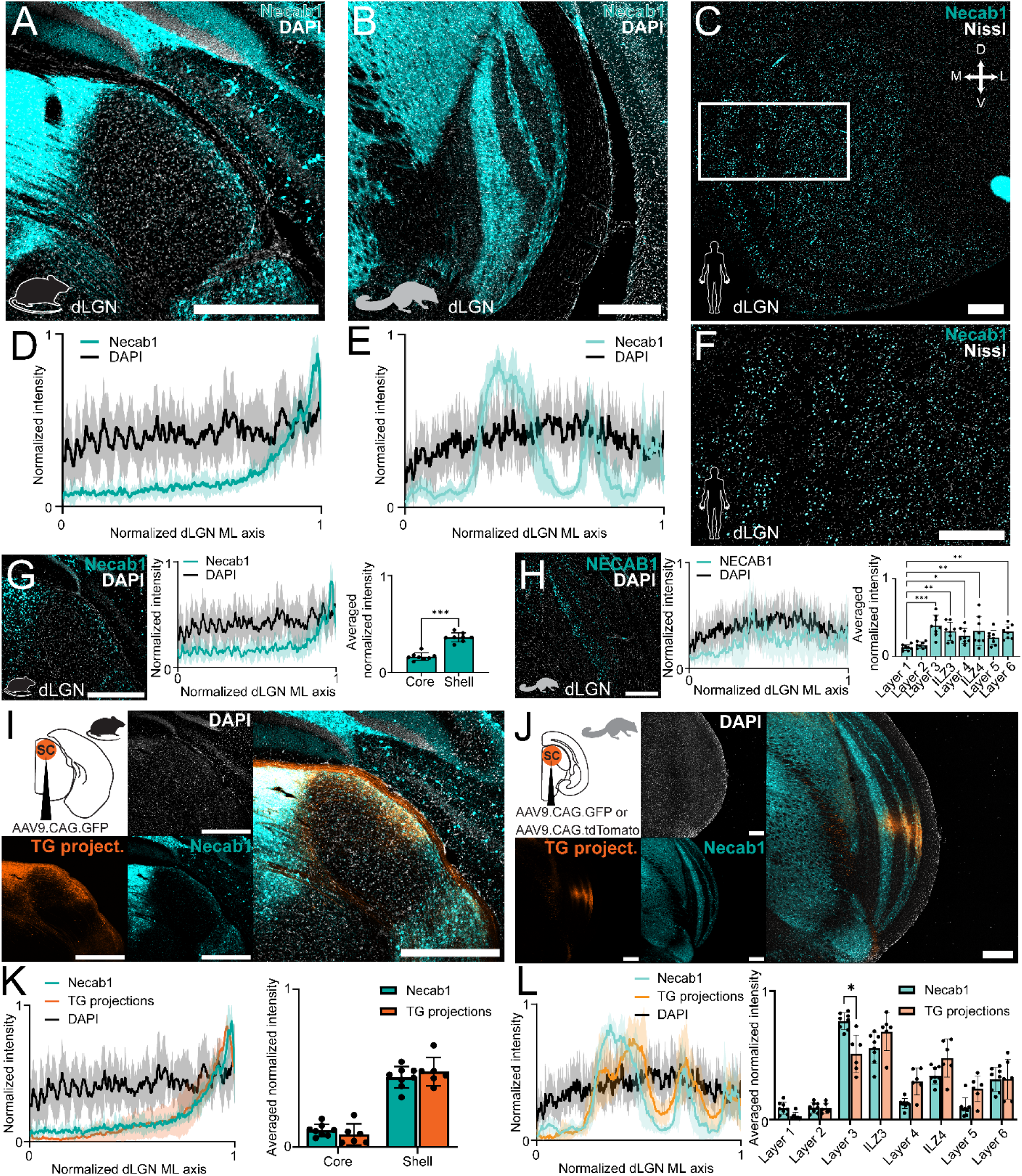
Colocalization of tectogeniculate projections and Necab1 expression using immunostaining and fluorescent in situ hybridization in mice, tree shrews, and humans. A and B. IHC in the mouse (A) and tree shrew (B) dLGN of Necab1 (cyan) with DAPI counterstain (white). C. IHC in the human dLGN of Necab1 (cyan) with Nissl counterstain (white). D and E. Normalized mean intensity profiles with shaded standard deviation of a 100 µm rectangular ROI spanning across the medial-lateral axis of the mouse (D) and tree shrew (E) dLGN (mouse: n = 6 animals, 8 sections; tree shrew: 4 animals, 8 sections). F. Photomicrograph of the white outlined region in (C). G. RNAscope FISH staining in mouse dLGN for Necab1 (cyan) with DAPI counterstain (white). Normalized mean intensity profile with shaded standard deviation of a 100 µm rectangular ROI spanning across the medial-lateral axis of the dLGN. Averaged normalized intensity binned for “core” and “shell” of the mouse dLGN using VGluT2-defined regions from Figure 1 (Mann-Whitney test; p ≤ 0.001 (***); core versus shell, p = 0.0002; n = 4 animals, 8 sections). H. Same as (G) but for the tree shrew. Averaged normalized intensity binned for layers and ILZs of the tree shrew dLGN using VGluT2-defined regions from Figure 1 (Kruskal-Wallis test with Dunn’s test for multiple comparison corrections comparing the mean rank of each column to the mean rank of Layer 1 with p ≤ 0.05 (*), ≤ 0.01 (**), and ≤ 0.001 (***); Kruskal-Wallis test, p < 0.0001; Dunn’s test comparing between layers 1 and 3, p = 0.0001; between layer 1 and ILZ3, p = 0.0029; between layers 1 and 4, p = 0.0336; between layer 1 and ILZ4, p = 0.0070, between layers 1 and 6, p = 0.0014; n = 2 animals, 8 sections). I. Anterograde tracing in the mouse with an injection of AAV9.CAG.GFP in the SC. Fibers from the SC (orange), with Necab1 IHC (cyan) and DAPI counterstain (white). J. Same as (I) but for the tree shrew. K. Normalized mean intensity profiles with shaded standard deviation of a 100 µm rectangular ROI spanning across the medial-lateral axis of the dLGN (Necab1: n = 6 animals, 8 sections; tectogeniculate projections: n = 6 animals, 6 sections). Averaged normalized intensity binned for “core” and “shell” of the mouse dLGN using VGluT2-defined regions from Figure 1 (Mann-Whitney test with Holm-Šídák correction comparing the mean rank of Necab1 and tectogeniculate projections data within the “core” and “shell” categories, in the core: p = 0.4047; in the shell: p = 0.8518). L. Same as (K) but for the tree shrew, averaged normalized intensity binned for layers and ILZs of the tree shrew dLGN using VGluT2-defined regions (Mann-Whitney test with Holm-Šídák correction comparing the mean rank of Necab1 and tectogeniculate projections data within each of the layers and ILZ categories with p ≤ 0.05 (*); in layer 1, p = 0.1318; in layer 2, p = 0.8858; in layer 3, p = 0.0367; in ILZ3, p = 0.1957; in layer 4, p = 0.1318; in ILZ4, p = 0.1957; in layer 5, p = 0.1957; in layer 6, p = 0.8858). All data presented as mean ± SD. All scale bars are 500 µm. Necab1 shows greater specificity for tectorecipient layers than Calbindin1

We next compared Necab1 with CB to determine whether Necab1 provides greater specificity for tectorecipient dLGN subdivisions than the classical K-pathway and K-pathway homolog marker. First, we compared the immunoreactivity of Necab1 and CB in the dLGN of both species. In the mouse dLGN, both Necab1 and CB show higher levels of protein expression in the shell than the core (Fig. 4A, top), and this is confirmed by comparing the intensity profiles (Fig. 4A, bottom left) as well as the binned intensity (Fig. 4A, bottom right). However, CB exhibits a higher expression than Necab1 throughout the mouse dLGN, indicating lower specificity (Fig. 4A, bottom right). In the tree shrew dLGN, there is a similar pattern. CB immunoreactivity is found throughout the dLGN while Necab1 shows high specificity for specific layers (Fig. 4B, top). Comparing the intensity profiles confirms this observation (Fig. 4B, bottom left) as do the binned intensities (Fig. 4B, bottom right). To assess the mRNA expression of Necab1 to that of Calb1, we performed RNAscope FISH in the mouse (Fig. 4C, top) and tree shrew dLGN (Fig. 4D, top). Comparing the intensity profiles of Necab1 and Calb1 in the mouse dLGN (Fig. 4C, bottom right) reveals that Necab1 mRNA expression is more restricted to the shell, whereas Calb1 mRNA expression is more broadly distributed throughout the dLGN (Fig. 4C, bottom left). In the tree shrew dLGN, NECAB1 shows similar mRNA expression to CALB1 (Fig. 4D, top and bottom right) and the difference in binned intensities is nonsignificant (Fig. 4D, bottom left). To visualize the relationship between Necab1 immunoreactivity and tectogeniculate projections, we used the same mouse and tree shrew dLGN sections shown for Necab1 and CB IHC experiments. These animals had received an anterograde tracer injection into the SC. Overlaying tracer-labeled tectogeniculate projections with Necab1 immunoreactivity revealed overlap within tectorecipient dLGN subdivisions in both species (mouse, Fig. 4E; tree shrew, Fig. 4F). Together, these results indicate that Necab1 shows greater specificity than CB for tectorecipient subdivisions of the mouse and tree shrew dLGN. We next addressed the neurotransmitter identity of Necab1 neurons in the mouse and the tree shrew and if expression of Necab1 correlates with direct SC inputs in mice.

**Figure 4.**
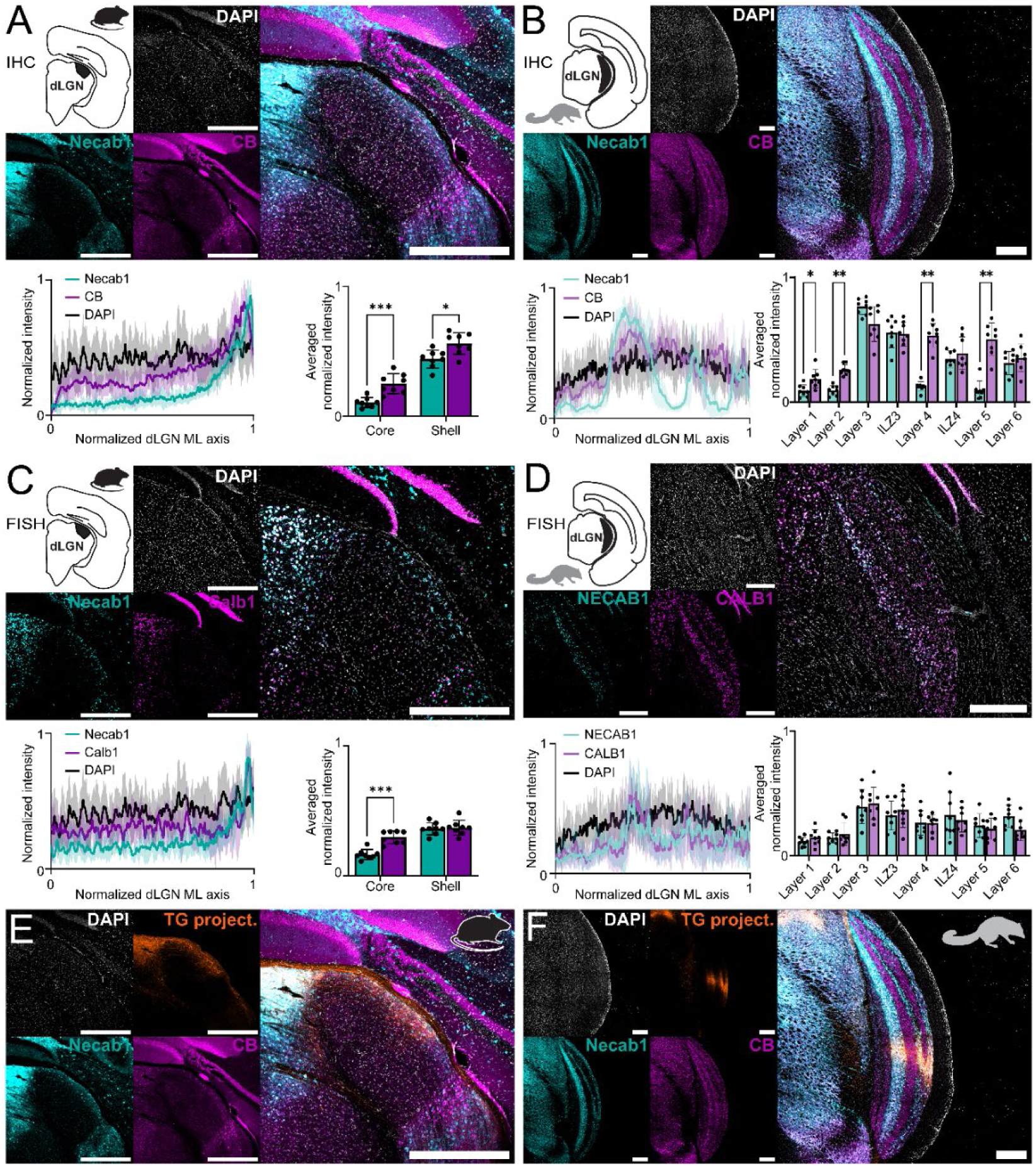
Necab1 shows higher specificity than CB for subdivisions of the mouse and tree shrew dLGN that receive tectogeniculate projections. A. IHC of Necab1 (cyan) and CB (magenta) in the mouse dLGN. Normalized mean intensity profile with shaded standard deviation of a 100 µm rectangular ROI spanning across the medial-lateral axis of the dLGN (n = 6 animals, 8 sections). Averaged normalized intensity binned for “core” and “shell” of the mouse dLGN using VGluT2-defined regions from Figure 1 (Mann-Whitney test with Holm-Šídák correction comparing the mean rank of Necab1 and CB data within the “core” and “shell” categories with p ≤ 0.05 (*) and ≤ 0.001 (***); in the core, p = 0.0006; in the shell, p = 0.0148). B. IHC of Necab1 (cyan) and CB (magenta) in the tree shrew dLGN. Normalized mean intensity profile with shaded standard deviation of a 100 µm rectangular ROI spanning across the medial-lateral axis of the dLGN (n = 4 animals, 8 sections). Averaged normalized intensity binned for layers and ILZs of the tree shrew dLGN using VGluT2-defined regions from Figure 1 (Mann-Whitney test with Holm-Šídák correction comparing the mean rank of Necab1 and CB data within each of the layers and ILZ categories with p ≤ 0.05 (*) and ≤ 0.01 (**); in layer 1, p = 0.0231; layer 2, p = 0.0012; layer 3, p = 0.1079; ILZ3, p = 0.9594; layer 4, p = 0.0012; ILZ4, p = 0.9594; layer 5, p = 0.0019; layer 6, p = 0.9554). C. Same as A but for RNAscope FISH (Mann-Whitney test with Holm-Šídák correction comparing the mRNA expression of Necab1 and Calb1 in the core, p = 0.0006; in the shell, p = 0.6454; n = 4 animals, 8 sections). D. Same as B but for FISH (Mann-Whitney test with Holm-Šídák correction comparing the mRNA expression of NECAB1 and CALB1 in layer 1, p = 0.9657; in layer 2, p = 0.9998; in layer 3, p = 0.9998; in ILZ3, p = 0.9980; in layer 4, p > 0.9999; in ILZ4, p = 0.9998; in layer 5, p = 0.9980; in layer 6, p = 0.1539; n = 2 animals, 8 sections). E and F. Photomicrographs of mouse (E) and tree shrew (F) dLGN with anterograde tracing from the SC (orange), with IHC of Necab1 (cyan) and CB (magenta) to illustrate the specificity of Necab1 for regions that receive tectogeniculate projections; same images from (A) and (B) but with tectogeniculate projections. All data presented as mean ± SD. All scale bars are 500 µm. Necab1+ cells are primarily glutamatergic and receive direct tectogeniculate inputs in mice.

To determine if Necab1+ cells are glutamatergic or GABAergic, we performed RNAscope FISH on mouse (Fig. 5A) and tree shrew (Fig. 5B) dLGN sections. Analysis in the mouse revealed that a majority of the Necab1+ cells are glutamatergic (Fig. 5C, left), with some cells expressing VGAT, which is seen in the normalized percentages (Fig. 5C, right; mean ± SD: VGLUT, 81.33 ± 5.84%; VGAT, 7.88 ± 2.57%). A similar relationship is seen in the tree shrew when comparing the raw counts (Fig. 5D, left) and normalized percentages (Fig. 5D, right; mean ± SD: VGLUT, 85.82 ± 3.60%; VGAT, 8.92 ± 3.36%), with a slight increase in the number of GABAergic cells compared to the mouse. Overall, the majority of Necab1+ neurons in the dLGN are glutamatergic.

**Figure 5.**
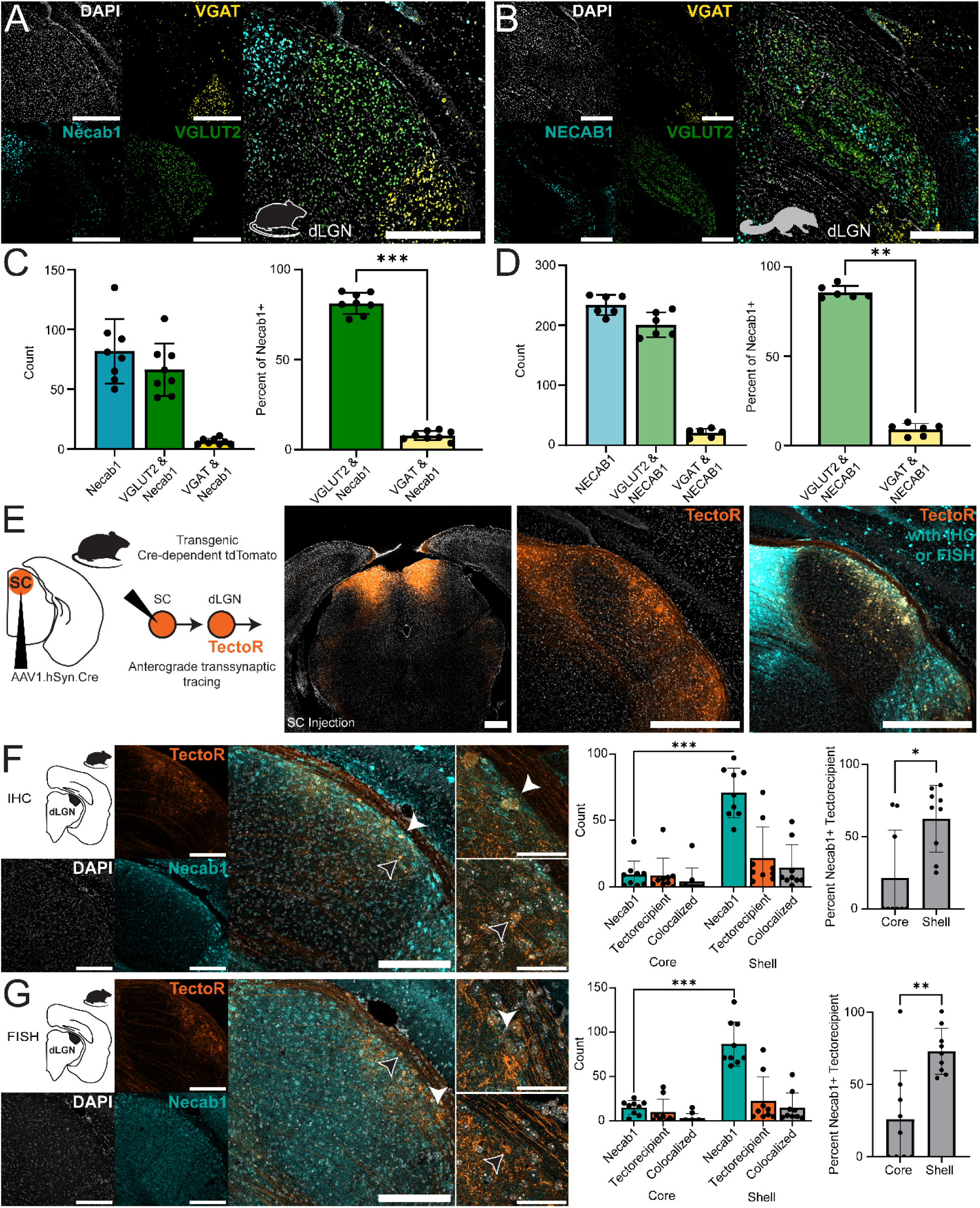
Immunostaining and fluorescent in situ hybridization of Necab1+ cells and colocalization with neurotransmitter and tectogeniculate inputs in the mouse and the tree shrew A. RNAscope FISH for VGAT (yellow), VGLUT2 (green), and Necab1 (cyan) in the mouse dLGN. B. Same as (A) but for the tree shrew dLGN. Scale bars for both (A) and (B) are 500µm. C. Left bar graph comparing the total count of Necab1+ cells (cyan), Necab1+ colocalized with VGLUT2+ (green), and Necab1+ colocalized with VGAT (yellow) for RNAscope FISH (VGAT+ and Necab1+: 50 total cells counted, mean ± SD: 6.25 ± 2.55 cells per section; VGLUT2+ and Necab1+: 531 total cells counted, mean ± SD: 66.38 ± 21.96 cells per section; total Necab1+: 654 total cells counted, mean ± SD: 81.75 ± 26.88 cells per section). Right bar graph comparing the percentage of Necab1+ colocalized with VGLUT2 (green) and Necab1+ colocalized with VGAT (yellow) (Mann-Whitney test; p ≤ 0.001 (***); Necab1+ cells that are also VGLUT2+ versus Necab1+ cells that are also VGAT+, p = 0.0002; n = 8 sections from 4 animals). D. Same as (C) but for the tree shrew (VGAT+ and NECAB1+: 123 total cells counted, mean ± SD: 20.50 ± 7.01 cells per section; VGLUT2+ and NECAB1+: 1205 total cells counted, mean ± SD: 200.83 ± 20.67 cells per section; total NECAB1+: 1402 total cells counted, mean ± SD: 233.67 ± 16.84 cells per section; NECAB1+ colocalized with VGLUT2, mean ± SD: 85.82 ± 3.60%; NECAB1+ colocalized with VGAT, mean ± SD: 8.92 ± 3.36%; Mann-Whitney test; p ≤ 0.01 (**); between the Necab1+ cells that are also VGLUT2+ versus Necab1+ cells that are also VGAT+, p = 0.0022; n = 6 sections from 2 animals). E. Schematic illustrating transsynaptic anterograde tracing combined with IHC or FISH. An injection of transsynaptic Cre in the SC of a Cre-dependent tdTomato reporter mouse line. Sections of the dLGN were collected and subsequently stained for Necab1 using either IHC or FISH. Scale bars for all photomicrographs in (E) are 500 µm. F. Photomicrographs of transsynaptic anterograde labeling (orange) and Necab1+ IHC (cyan) counterstained using DAPI (white). White arrow illustrates a positive cell, and the black arrow illustrates a negative cell. Scale bars for the representative staining photomicrograph are 200 µm while the scale bars for the higher magnification photomicrographs are 50 µm. Left bar graph comparing the count of all Necab1+ cells (cyan), all tectorecipient cells (orange), and colocalized (grey) (Mann-Whitney test comparing the count of Necab1+ cells in the core to that in the shell with p ≤ 0.001 (***); p = 0.0001; total Necab1+: 719 cells counted, mean ± SD: 79.89 ± 24.34 cells per section; total tectorecipient: 270 cells counted, mean ± SD: 30.00 ± 35.23 cells per section; Necab1+ and tectorecipient (colocalized): 167 cells counted, mean ± SD: 18.56 ± 26.38 cells per section). Right bar graph comparing the percentage of Necab1+ tectorecipient cells in the shell and the core (Mann-Whitney test, p ≤ 0.05 (*); p = 0.0112; n = 9 sections from 3 animals). G. Same as (F) but for Molecular Instruments HCR mRNA-FISH (Mann-Whitney test comparing between the number of Necab1+ cells in the core and the shell, p < 0.0001 (***); p = 0.0001; Mann-Whitney test comparing between the percentage of Necab1+ tectorecipient cells in the core and the shell, p < 0.01 (**); p = 0.0028; n = 9 sections from 3 animals; total Necab1+: 914 cells counted, mean ± SD: 101.56 ± 29.62 cells per section; total tectorecipient: 289 cells counted, mean ± SD: 32.11 ± 41.30 cells per section; Necab1+ and tectorecipient (colocalized): 160 cells counted, mean ± SD: 17.78 ± 21.35 cells per section). All data presented as mean ± SD.

To determine if the dLGN cells that receive tectogeniculate projections (tectorecipient cells, tectoR) express Necab1, we performed anterograde transsynaptic viral tracing using a transgenic mouse line expressing Cre-dependent tdTomato and an anterograde transsynaptic Cre virus and combined this with IHC and FISH experiments (Fig. 5E). Raw counts of the IHC experiment revealed that a majority of tectorecipient cells express Necab1 (Fig. 5F, white arrow), with some cells lacking expression (Fig. 5F, black arrow), and a majority of the Necab1+ cells located in the shell, which was also true for tectorecipient cells (Fig. 5F, raw count bar graph; Fig. 5F, percentage bar graph; *shell tectorecipient Necab1+ mean ± SD: 62.30 ± 23.06%; core tectorecipient Necab1+: 21.50 ± 32.86%*). The same relationship was seen in the FISH experiment (Fig. 5G, white arrow to indicate a positive cell; black arrow to indicate a negative cell). A majority of the Necab1+ cells were located in the shell, including the tecorecipient cells (Fig. 5G, raw count bar graph; Fig, 5G, percentage bar graph; *shell tectorecipient Necab1+ mean ± SD: 72.96 ± 15.80%; core tectorecipient Necab1+: 26.03 ± 33.60%;*). These results suggest that in the shell of the dLGN of the mouse, the majority of tectorecipient cells express Necab1.

### Tectogeniculate neurons are mostly glutamatergic in mice

To begin to uncover the molecular identity of the SC cells that project to the dLGN (tectogeniculate cells), we performed an injection of a retrograde tracer in the dLGN of the mouse, collected sections of the SC, and stained those sections for VGLUT2 and VGAT using Molecular Instruments HCR mRNA-FISH (Fig. 6A). The mRNA-FISH experiment showed a population of retrograde labeled SC cells that were positive for VGLUT2 (Fig. 6B, green, white arrow; Fig. 6E, white arrow) and another population that was positive for VGAT (Fig. 6B, yellow, black arrow; Fig. 6D, black arrow). Quantification of these cells shows a majority of tectogeniculate cells expressing VGLUT2 mRNA, however we could also identify a small proportion of VGAT+ neurons (Fig. 6C, left). Comparing the percentage of VGLUT2+ and VGAT+ tectogeniculate cells also confirmed that a majority of the cells are VGLUT2+ (Fig. 6C, right; percentage of VGLUT2+ tectogeniculate cells, mean ± SD: 81.48 ± 7.14%; percentage of VGAT+ tectogeniculate cells, mean ± SD: 5.39 ± 3.71%). Not every tectogeniculate cell was positive for either VGLUT2 or VGAT mRNA, as some cells colocalized both or neither. These cells were not included in the analysis (20 cells were excluded from 137 total retrograde labeled cells). The results of mRNA-FISH experiments as well as the seemingly various morphologies displayed from tectogeniculate cells (Fig. 6D and 6E) provide evidence for multiple subpopulations of tectogeniculate cells in the mouse SC.

**Figure 6.**
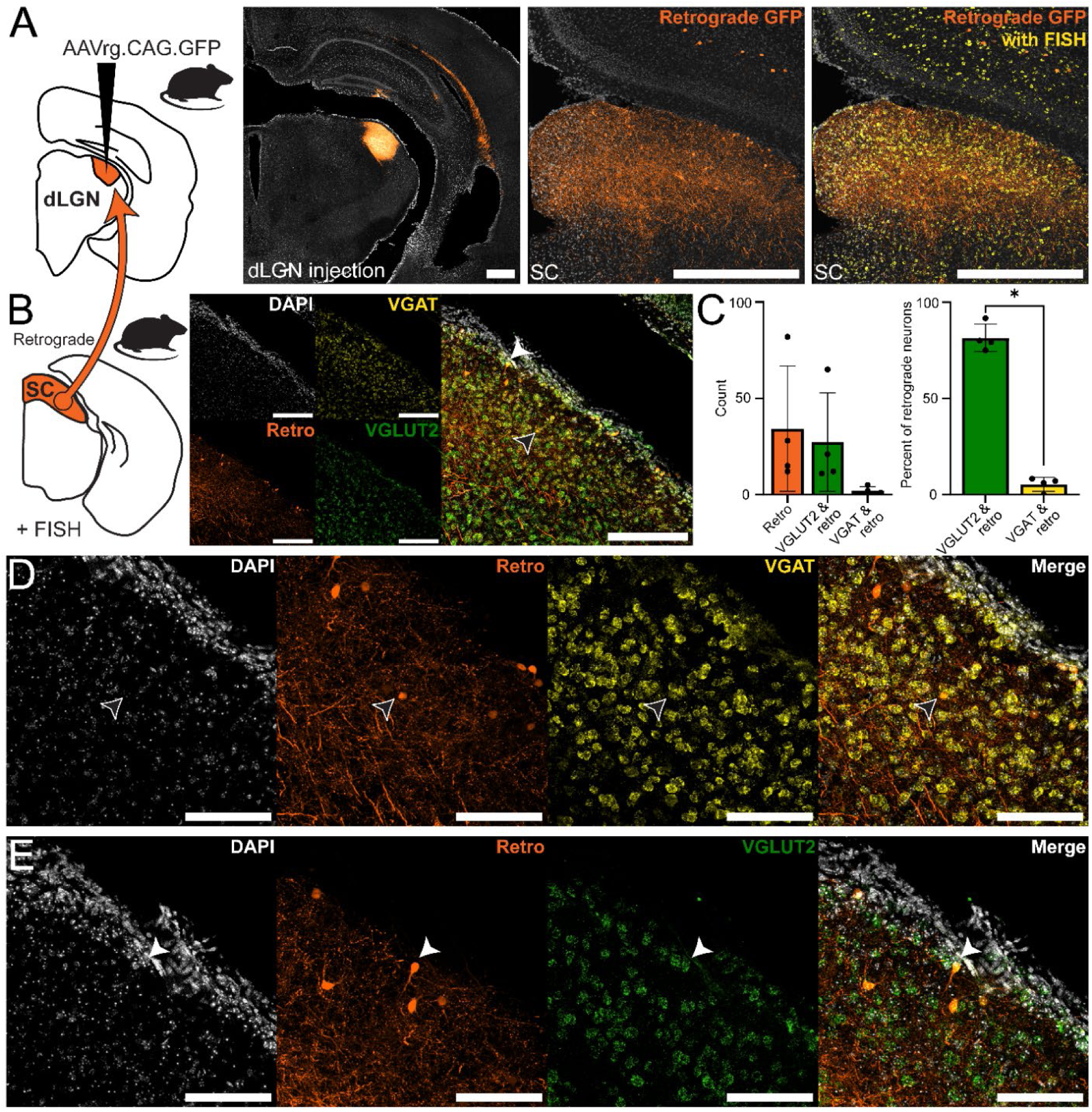
Retrograde tracing of tectogeniculate projections and identification of neurotransmitter expression by fluorescent in situ hybridization in the mouse A. Schematic to illustrate experiment protocol. An injection of a retrograde tracer was performed in the mouse dLGN and sections of the SC were collected and subsequently stained using Molecular Instruments HCR mRNA-FISH. All scale bars for (A) are 500 µm. B. Results of retrograde tracing experiment (labeled SC cells in orange) combined with FISH staining for VGAT (yellow) and VGLUT2 (green) counterstained using DAPI (white). Black arrow indicates a cell positive for VGAT and white arrow indicates a cell positive for VGLUT2. Scale bars for (B) are 200 µm. C. Left bar graph comparing the raw counts from the retrograde tracing experiment for retrograde labeled SC cells (orange), retrograde labeled and VGLUT2 (green), and retrograde labeled and VGAT (yellow). Right bar graph comparing the percentage of retrograde labeled cells that are colocalized with VGLUT2 (green) and VGAT (yellow) (Mann-Whitney test, p ≤ 0.05 (*); p = 0.0286; n = 4 animals across 11 sections with 2-4 sections per animal; data presented are the counts for each animal across all sections sampled within that animal; retrograde labeled: 137 total cells, mean ± SD: 34.25 ± 32.58 cells per animal; retrograde labeled and VGLUT2+: 109 total cells, mean ± SD: 27.25 ± 25.57 cells per animal; retrograde labeled and VGAT+: 8 total cells, mean ± SD: 2.00 ± 2.16 cells per animal). D and E. Photomicrographs indicating a retrograde labeled SC cell (orange) positive for VGAT (D, yellow, black arrow) and VGLUT2 (E, green, white arrow). Scale bars for (D and E) are 100 µm. All data presented as mean ± SD.

## Discussion

Leveraging the wealth of information acquired in mice at the microcircuit and cell-type level to organisms with distinct sensory specialization is a required step to investigate physiology and behavior linked to sensory processing. Tree shrews are phylogenetically close to primates and display high visual function, thus enabling studies of visually guided behavior. However, it remains unknown how cell-types and circuits are conserved between the mouse and the tree shrew. Here, we compare the anatomical organization and connectivity of mouse and tree shrew dLGN, revealing key differences and similarities. Our goal was to identify a conserved genetic access point to investigate interactions between the SC and dLGN, two evolutionarily distinct visual centers. Our main hypothesis was that despite anatomical differences, a conserved molecular marker could be identified. We proceeded with the characterization of anatomical subdivisions in the dLGN and examined the expression pattern of molecular markers.

### Anatomical differences in the dLGN of the mouse and the tree shrew

The mouse dLGN lacks clear lamination, however, response properties as well as innervation are graded along the medial-lateral axis, forming the core and the shell (Harting et al., 1991; Gale and Murphy, 2014; Bickford et al., 2015; Guido, 2018; Fei et al., 2025). Here, we developed a quantitative approach to delineate the shell from the core using VGluT2 immunoreactivity and confirmed that tectogeniculate inputs are restricted to the shell in mice. In the tree shrew dLGN, the reported organization of tectogeniculate projections has been more variable. Previous studies have noted tectogeniculate projections in a region medial to layer 1, in layers 3 and 6, and the ILZ between layers 4 and 5 (Fitzpatrick et al., 1980). Others have reported projections to the ILZs between layers 2 and 3, between layers 3 and 4, and between layers 4 and 5, in addition to layers 3 and 6 (Harting et al., 1991). Across studies, however, the most consistent finding has been SC input to layers 3 and 6 (Diamond et al., 1991; Sciaccotta et al., 2025). Our results similarly revealed SC projections to layer 3 and layer 6, as well as the ILZs between layers 3 and 4 and between layers 4 and 5. It is important to note that the distribution and intensity of these projections depends on the location and size of the SC injection site as well as the anterior-posterior position within the dLGN. This is a phenomenon that has been noted by others as well (Fitzpatrick et al., 1980; Harting et al., 1991).

### Proportion of GABAergic and glutamatergic neurons in the mouse and tree shrew dLGN

Both the mouse and tree shrew dLGN contained a higher proportion of glutamatergic than GABAergic cells. However, the tree shrew dLGN contained a greater proportion of VGAT+ cells than the mouse dLGN, suggesting species-specific differences in inhibitory interneurons. This difference can also be found in the broader context of thalamic evolution, as the prevalence of local inhibitory neurons varies substantially across mammalian species (Winer and Larue, 1996). The greater relative proportion of VGAT+ cells in the dLGN of the highly visual tree shrew is in line with these observations and suggests that subcortical inhibition is associated with sensory specialization.

### Leveraging molecular markers to understand visual thalamus organization across species

In adult macaque monkeys, PV is expressed in both the M and P layers and largely absent from the ILZs (Yan et al., 1996). In contrast, CB is expressed in the ILZs, which are termed the K layers in primates (Murray et al., 2008), while CR is found almost exclusively in the medial-most ILZ known as the “S” layer (Yan et al., 1996). Within our histological dataset, differences in expression of protein versus mRNA for the same biomarker were observed. For example, CR immunoreactivity is present in the mouse dLGN, however no CR mRNA was detected in this region. The CR immunoreactivity can be attributed to projections from retina and SC, both of which express CR locally (Yang and Jeon, 1998; Soares et al., 2001; Huberman et al., 2008).

In tree shrews, PV expression was reported in layers 1, 2, 4, and 5 while CB is present in layers 4, 5, and 6 and CR can be found across all layers (Zhang et al., 2025). We also find heightened expression of PV in layers 2 and 4 in the tree shrew dLGN. The classical molecular markers examined in this study, PV, CB, and CR are all Ca^2+^-binding proteins, but their physiological role may differ across brain regions. For example, PV is associated with fast-spiking inhibitory interneurons within the cortex (Baimbridge et al., 1992), but also excitatory relay cells of the primate dLGN (Jones and Hendry, 1989; Ma et al., 2023). Similarly, CB has previously been used as a marker of K layers in the primate dLGN (Jones and Hendry, 1989; Diamond et al., 1993; Hendry and Yoshioka, 1994; Goodchild and Martin, 1998; Hendry and Reid, 2000), but its expression is not restricted to tectorecipient regions in all species as shown in this study.

Our findings identify Necab1 as a conserved molecular marker of tectorecipient subdivisions of the dLGN in both the mouse and the tree shrew, suggesting that molecular features can be conserved across species despite differences in anatomical organization. Necab1 belongs to a distinct family of neuronal Ca^2+^-binding proteins (NECABs) characterized by unique Ca^2+^-binding domains for eukaryotic cells. Necab1 has been reported in discrete neuronal populations, including layer 4 pyramidal neurons of the mouse cerebral cortex and inhibitory interneurons in the hilus of the mouse hippocampus (Sugita et al., 2002). In the context of this study, Necab1 shows greater selectivity for dLGN subdivisions that receive tectogeniculate inputs in both the mouse and tree shrew when compared to CB. We also show that Necab1 neurons in the mouse receive direct inputs from tectogeniculate projections. We could not replicate these findings in tree shrews due to the lack of transgenic lines. However, by conducting our experiments in parallel with two distinct species, we have confirmed that the molecular definition of cell-types can also depend on its connectivity and neurotransmitter properties, an essential step to understanding the implementation of neuronal computations. Indeed, we confirmed that Necab1 neurons are mostly glutamatergic both in the mouse and the tree shrew. This is in line with previous studies suggesting that tectogeniculate projections are driver-like and could substantially modulate visual information within the dLGN (Bickford et al., 2015). Interestingly, the presence of Necab1 extends to the human dLGN, leaving open intriguing possibilities regarding molecular marker conservation.

### Heterogeneity of GABAergic and glutamatergic tectogeniculate projections

Our retrograde tracing and mRNA-FISH results indicate that tectogeniculate neurons are primarily glutamatergic, although a smaller VGAT+ population is also present. These data are consistent with previous studies showing that both glutamatergic and GABAergic SC neurons project to the mouse dLGN, although reported proportions vary across studies (Gale and Murphy, 2014; Bickford et al., 2015; Li et al., 2023). Together with our findings, these studies suggest that tectogeniculate projection neurons are comprised of multiple molecular subtypes. Consistent with this idea, transcriptomic-based analyses have proposed RorB as a broad marker for SC neurons projecting to visual thalamic nuclei (Byun et al., 2019), yet more recent evidence indicates that RorB-expressing cells in the mouse SC include both glutamatergic and GABAergic populations (Liu et al., 2023). These findings indicate that this highly conserved tectogeniculate pathway should be considered a heterogeneous pathway, with distinct cell-types used to convey visual information. The heterogeneity of the tectogeniculate projection neurons extends beyond molecular identity, as these neurons also appear to be morphologically and functionally diverse (Fei et al., 2025; Sciaccotta et al., 2025).

### What is a cell type and how can these definitions inform understanding of the evolution of sensory processing?

Early efforts to classify neuronal cell types relied heavily on morphology (Ramón Y Cajal, 1909). Modern approaches make it possible to classify cell types with molecular and functional definitions (Bugeon et al., 2022; Liu et al., 2023). However, these approaches have also made clear that a cell type is not defined by a single feature, but rather a convergence of properties including gene expression, morphology, connectivity, and function. This is especially important when comparing sensory circuits across species, as a molecular marker identified in one species might not be conserved in another. For example, transcriptomic analyses suggest that wide-field vertical cells in the mouse SC are best defined using the gene NPNT. However, NPNT does not selectively label the same population of cells in the tree shrew SC. Instead, Cbln2 provides a more conserved marker across both species (Liu et al., 2023; Kipcak and Erisir, 2026). In this study, Necab1 serves this role for tectorecipient regions of the mouse and tree shrew dLGN. Necab1 not only identifies these anatomical regions, but also provides a genetic access point for isolating tectogeniculate projections across species (Liu et al., 2023; Kipcak and Erisir, 2026). Together, our comparative framework between the mouse and the tree shrew demonstrates how integrating molecular identity, anatomy, and connectivity can reveal conserved and species-specific features of visual thalamic organization.

The authors declare no competing financial interests

## Acknowledgements

The authors acknowledge the tissue donors and their families and thank the NIH NeuroBioBank and the University of Michigan Brain Bank for providing the specimens used in this study. University of Michigan Brain Bank specimens were, in part, obtained from the NIH Brain & Tissue Repository-California, Human Brain & Spinal Fluid Resource Center, VA West Los Angeles, Medical Center, Los Angeles, California, which is supported in part by the NIH and the US Department of Veterans Affairs.

---

**Supplemental Figure 1.**
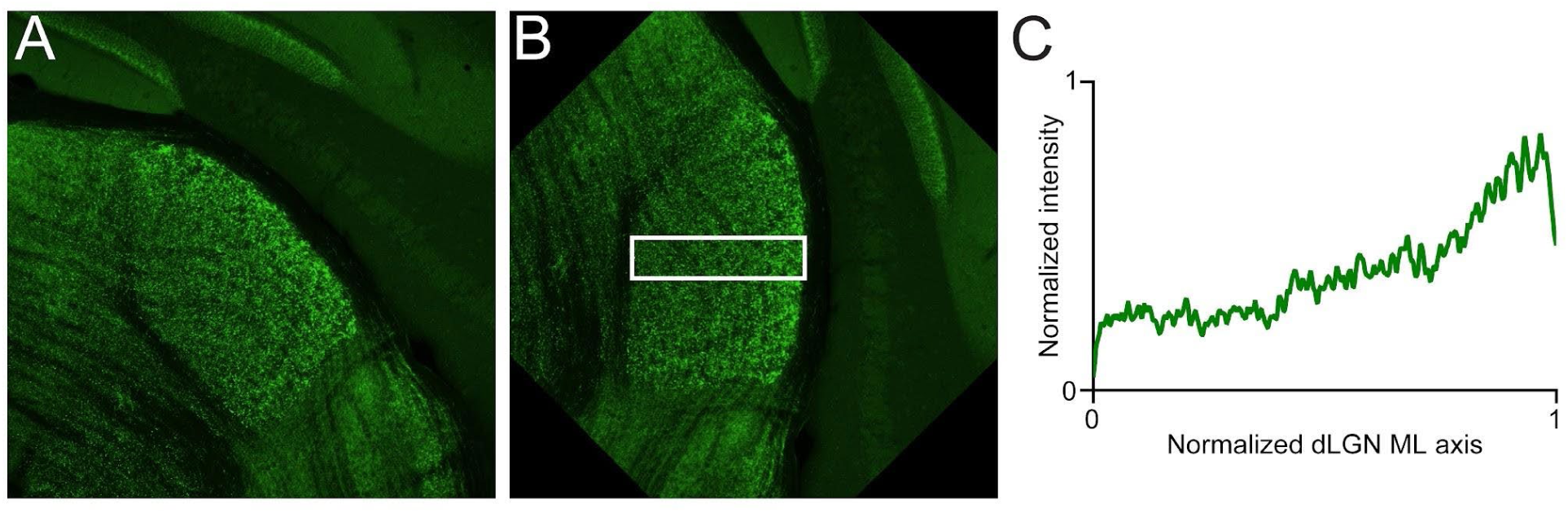
Example workflow for intensity profile collection. **A.** Maximum intensity projections were generated from 10x Z-stacks, shown here is an example of VGluT2 IHC staining (green). **B.** Maximum intensity projections were rotated so that the lateral-most edge of the dLGN is perpendicular to the lower edge boundary. A 100µm rectangular region of interest was drawn over the rotated image (white rectangle). **C.** An intensity profile of the rectangular region of interest was collected for further analysis.

**Supplemental Figure 2.**
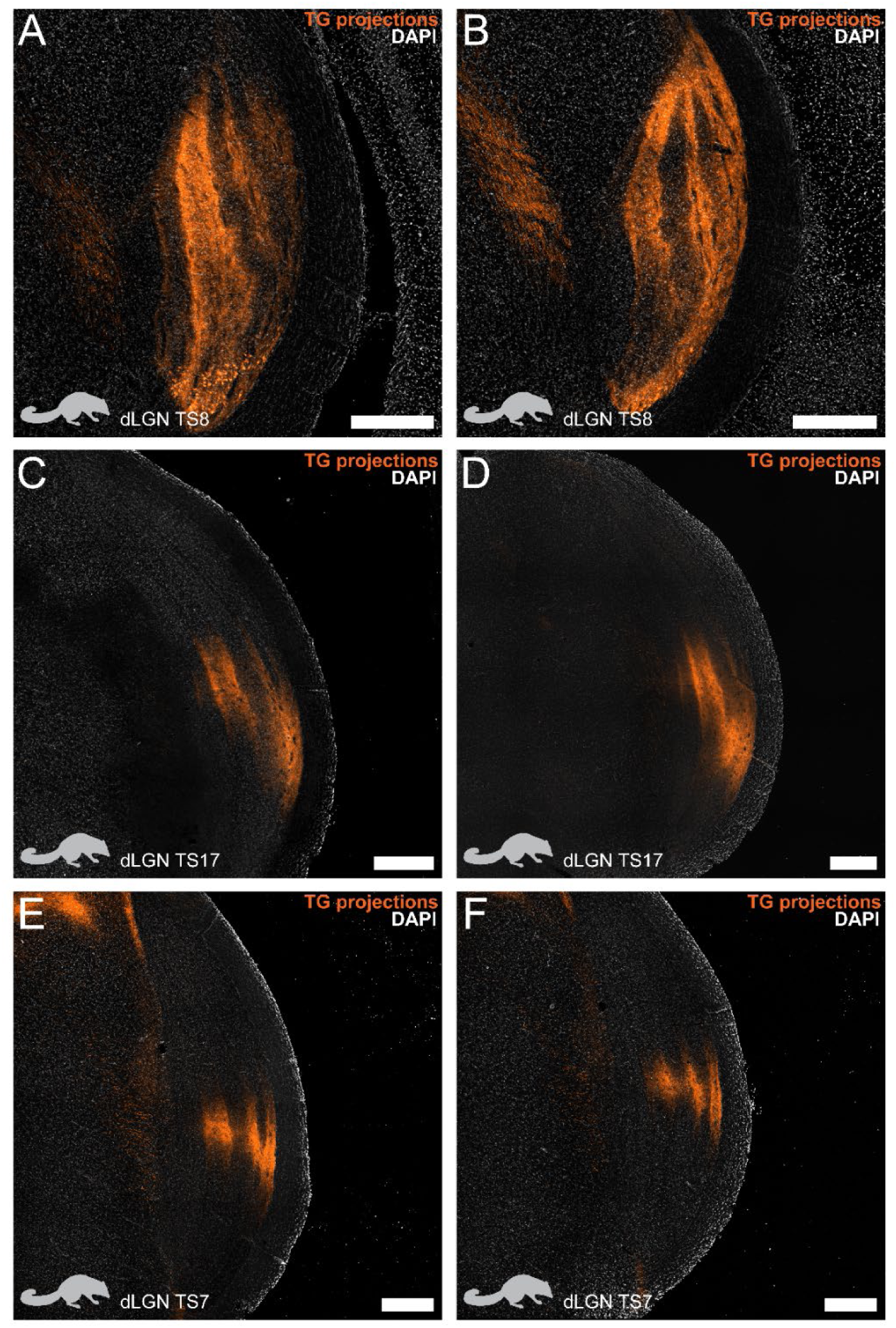
Tectogeniculate projections in the dLGN of the tree shrew. **A-F.** Tectogeniculate projections (orange) in the dLGN of the tree shrew counterstained with DAPI (white) to illustrate the variability present throughout the anterior-posterior axis. (**A**) and (**B**) are sections from TS8, (**C**) and (**D**) are sections from TS17, (**E**) and (**F**) are from TS7. All scale bars 500 µm.

**Supplemental Figure 3.**
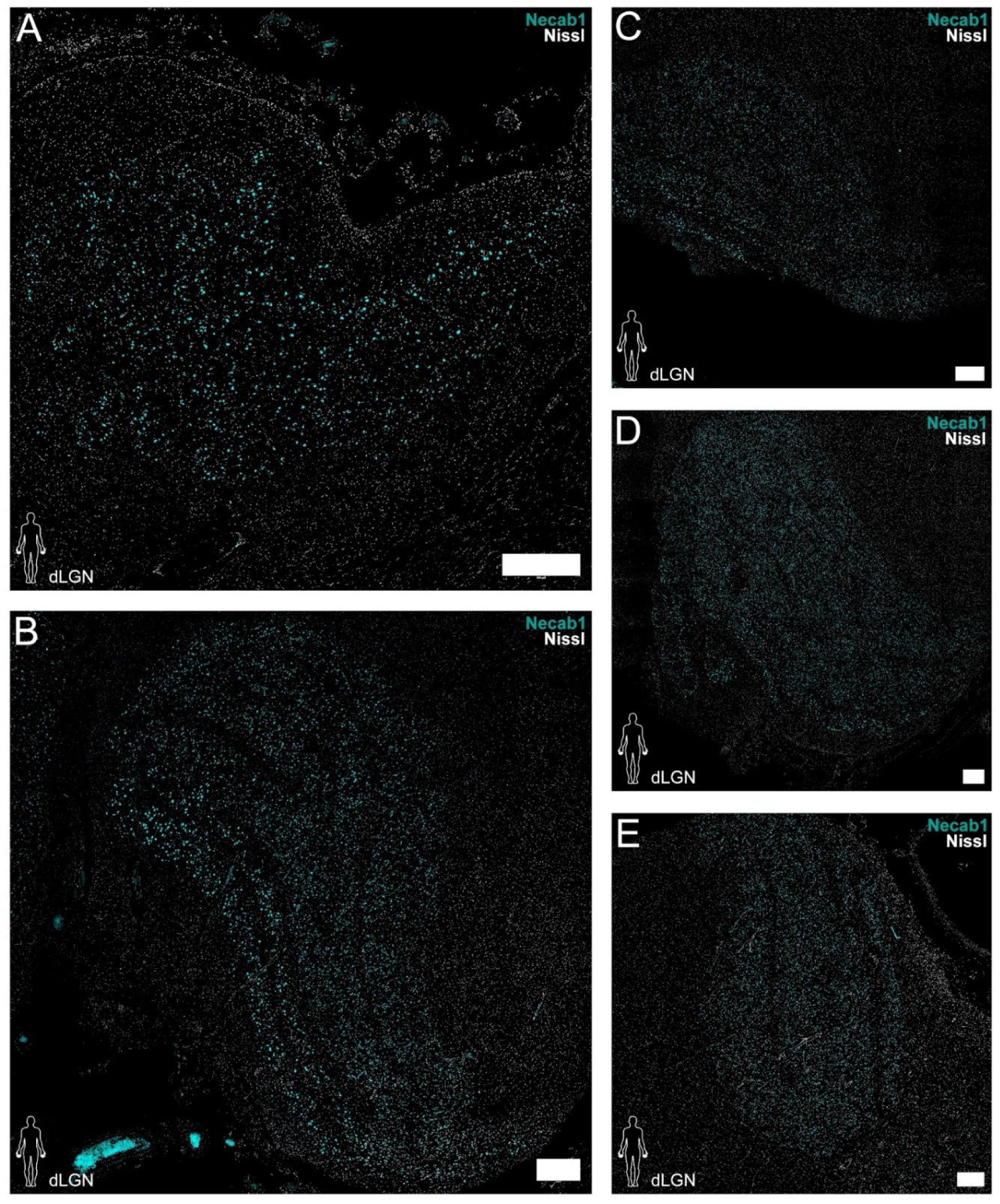
IHC staining of human dLGN for Necab1. **A-E.** IHC staining of human dLGN for Neacb1 (cyan) counterstained with Nissl staining (grey) across 5 different donors. Photomicrographs illustrate the specificity of Necab1 for the dLGN as well as the variability of the layers and size within the human dLGN. All scale bars are 500 µm.

**Supplemental Figure 4.**
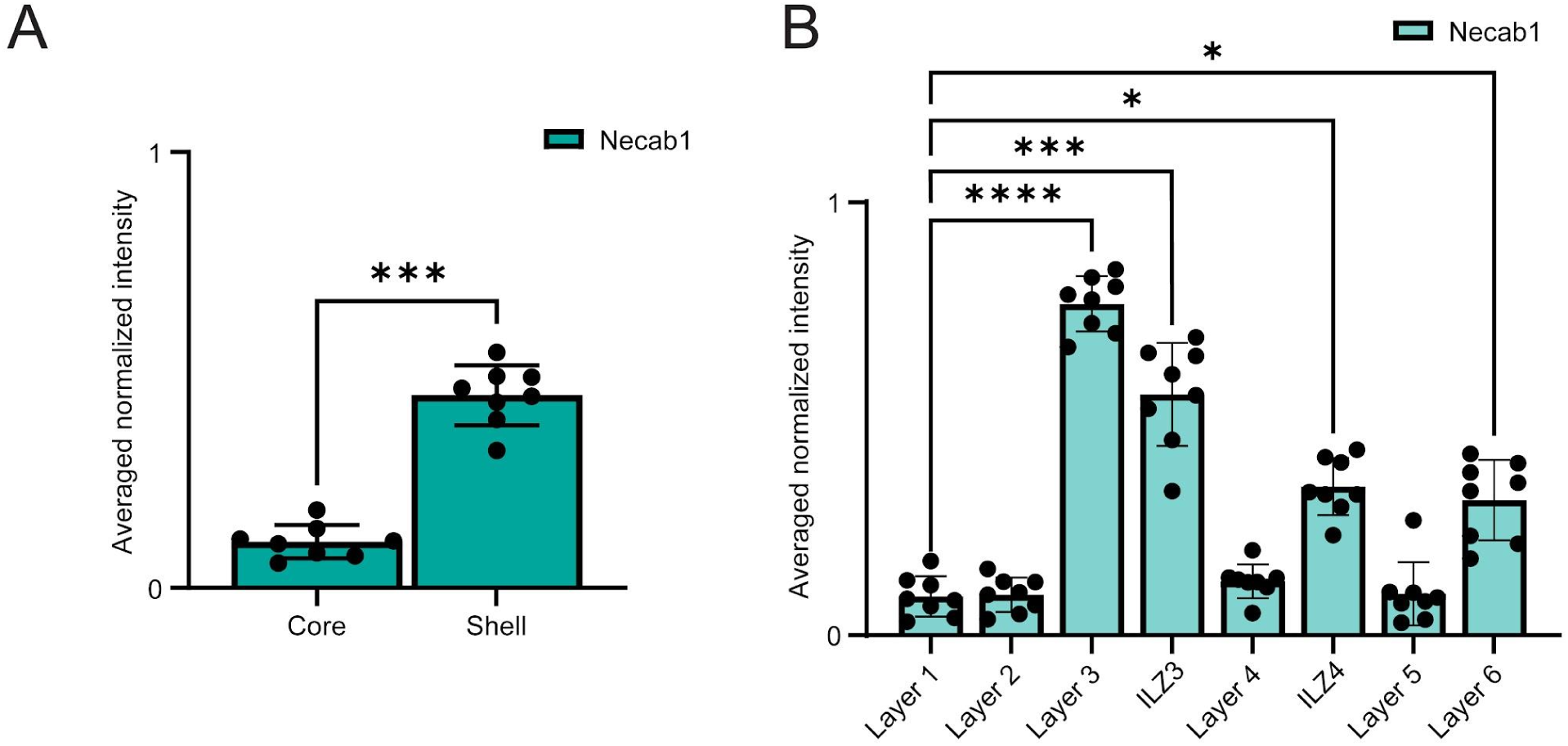
Binned Necab1 IHC intensity profiles. **A.** Averaged normalized intensity binned for “core” and “shell” of the mouse dLGN Necab1 IHC staining using VGluT2-defined regions from **Figure 1**, Mann-Whitney test; p ≤ 0.001 (***) (n = 8 sections from 6 animals). **B.** Same as (**A**) but for the tree shrew. Averaged normalized intensity binned for layers and ILZs of the tree shrew dLGN Necab1 IHC staining using VGluT2-defined regions from **Figure 1**, Kruskal-Wallis test with Dunn’s test for multiple comparison corrections comparing the mean rank of each column to the mean rank of Layer 1 with p ≤ 0.05 (*), ≤ 0.01 (**), ≤ 0.001 (***), and ≤ 0.0001 (****) (n = 8 sections from 4 animals).

## Notes

https://doi.org/10.7302/y6cm-h505

### Competing Interest Statement

The authors have declared no competing interest.

### Funding

- NIH National Eye Institute: R00EY031783

## References

Baimbridge KG, Celio MR, Rogers JH (1992) Calcium-binding proteins in the nervous system. Trends Neurosci 15:303–308.

Bakken TE et al. (2021) Single-cell and single-nucleus RNA-seq uncovers shared and distinct axes of variation in dorsal LGN neurons in mice, non-human primates, and humans. eLife 10:e64875.

Balaram P, Isaamullah M, Petry HM, Bickford ME, Kaas JH (2015) Distributions of vesicular glutamate transporters 1 and 2 in the visual system of tree shrews (*Tupaia belangeri*): VGLUT1 and VGLUT2 In Tree Shrew Visual Areas. J Comp Neurol 523:1792–1808.

Basso MA, Bickford ME, Cang J (2021) Unraveling circuits of visual perception and cognition through the superior colliculus. Neuron 109:918–937.

Bickford ME, Zhou N, Krahe TE, Govindaiah G, Guido W (2015) Retinal and Tectal “Driver-Like” Inputs Converge in the Shell of the Mouse Dorsal Lateral Geniculate Nucleus. Journal of Neuroscience 35:10523–10534.

Bugeon S, Duffield J, Dipoppa M, Ritoux A, Prankerd I, Nicoloutsopoulos D, Orme D, Shinn M, Peng H, Forrest H, Viduolyte A, Reddy CB, Isogai Y, Carandini M, Harris KD (2022) A transcriptomic axis predicts state modulation of cortical interneurons. Nature 607:330–338.

Byun H, Lee H-L, Liu H, Forrest D, Rudenko A, Kim I-J (2019) Rorβ regulates selective axon-target innervation in the mammalian midbrain. Development 146:dev171926.

Cang J, Savier E, Barchini J, Liu X (2018) Visual Function, Organization, and Development of the Mouse Superior Colliculus. Annu Rev Vis Sci 4:239–262.

Casagrande VA (1994) A third parallel visual pathway to primate area VI. Trends in Neurosciences. 17:305–310.

Cruz-Martín A, El-Danaf RN, Osakada F, Sriram B, Dhande OS, Nguyen PL, Callaway EM, Ghosh A, Huberman AD (2014) A dedicated circuit links direction-selective retinal ganglion cells to the primary visual cortex. Nature 507:358–361.

Diamond IT, Conley M, Fitzpatrick D, Raczkowski D (1991) Evidence for separate pathways within the tecto-geniculate projection in the tree shrew. Proceedings of the National Academy of Sciences 88:1315–1319.

Diamond IT, Fitzpatrick D, Schmechel D (1993) Calcium binding proteins distinguish large and small cells of the ventral posterior and lateral geniculate nuclei of the prosimian galago and the tree shrew (Tupaia belangeri). Proceedings of the National Academy of Sciences 90:1425–1429.

Fei Y, Luh MY, Ontiri A, Ghauri D, Hu W, Liang L (2025) Coordination of distinct sources of excitatory inputs enhances motion selectivity in the mouse visual thalamus. Neuron 113:3441–3457.e13.

Fitzpatrick D, Carey RG, Diamond IT (1980) The projection of the superior colliculus upon the lateral geniculate body in tupaia glis and galago senegalensis. Brain Research 194:494–499.

Fremeau RT, Voglmaier S, Seal RP, Edwards RH (2004) VGLUTs define subsets of excitatory neurons and suggest novel roles for glutamate. Trends in Neurosciences 27:98–103.

Gale SD, Murphy GJ (2014) Distinct Representation and Distribution of Visual Information by Specific Cell Types in Mouse Superficial Superior Colliculus. Journal of Neuroscience 34:13458–13471.

Goodchild AK, Martin PR (1998) The distribution of calcium-binding proteins in the lateral geniculate nucleus and visual cortex of a New World monkey, the marmoset, *Callithrix jacchus*. Vis Neurosci 15:625–642.

Grubb MS, Thompson ID (2004) Biochemical and anatomical subdivision of the dorsal lateral geniculate nucleus in normal mice and in mice lacking the β2 subunit of the nicotinic acetylcholine receptor. Vision Research 44:3365–3376.

Guido W (2018) Development, form, and function of the mouse visual thalamus. Journal of Neurophysiology 120:211–225.

Harting JK, Huerta MF, Hashikawa T, van Lieshout DP (1991) Projection of the mammalian superior colliculus upon the dorsal lateral geniculate nucleus: Organization of tectogeniculate pathways in nineteen species. The Journal of Comparative Neurology 304:275–306.

Hendry SHC, Reid RC (2000) The Koniocellular Pathway in Primate Vision. Annu Rev Neurosci 23:127–153.

Hendry SHC, Yoshioka T (1994) A Neurochemically Distinct Third Channel in the Macaque Dorsal Lateral Geniculate Nucleus. Science 264:575–577.

Huberman AD, Manu M, Koch SM, Susman MW, Lutz AB, Ullian EM, Baccus SA, Barres BA (2008) Architecture and Activity-Mediated Refinement of Axonal Projections from a Mosaic of Genetically Identified Retinal Ganglion Cells. Neuron 59:425–438.

Johnson JK, Casagrande VA (1995) Distribution of calcium-binding proteins within the parallel visual pathways of a primate (*Galago crassicaudatus*). J of Comparative Neurology 356:238–260.

Jones EG, Hendry SHC (1989) Differential Calcium Binding Protein Immunoreactivity Distinguishes Classes of Relay Neurons in Monkey Thalamic Nuclei. Eur J of Neuroscience 1:222–246.

Kipcak A, Erisir A (2026) CBLN2 promoter enables genetic access to wide-field neurons of the tree shrew superior colliculus. Cell Rep Methods 6:101309.

Lachica EA, Casagrande VA (1993) The morphology of collicular and retinal axons ending on small relay (W-like) cells of the primate lateral geniculate nucleus. Vis Neurosci 10:403–418.

Li C, Kühn NK, Alkislar I, Sans-Dublanc A, Zemmouri F, Paesmans S, Calzoni A, Ooms F, Reinhard K, Farrow K (2023) Pathway-specific inputs to the superior colliculus support flexible responses to visual threat. Sci Adv 9:eade3874.

Liang L, Chen C (2020) Organization, Function, and Development of the Mouse Retinogeniculate Synapse. Annu Rev Vis Sci 6:261–285.

Liu Y, McDaniel JA, Chen C, Yang L, Kipcak A, Savier EL, Erisir A, Cang J, Campbell JN (2025) Co-Conservation of synaptic gene expression and circuitry in collicular neurons. Nat Commun 16:9146.

Liu Y, Savier EL, DePiero VJ, Chen C, Schwalbe DC, Abraham-Fan R-J, Chen H, Campbell JN, Cang J (2023) Mapping visual functions onto molecular cell types in the mouse superior colliculus. Neuron 111:1876–1886.e5.

Ma G, Worthy KH, Liu C, Rosa MGP, Atapour N (2023) Parvalbumin as a neurochemical marker of the primate optic radiation. iScience 26:106608.

Martin PR, Solomon SG (2019) The koniocellular whiteboard. J Comp Neurol 527:505–507.

May PJ (2006) The mammalian superior colliculus: laminar structure and connections. In: Progress in Brain Research, pp 321–378. Elsevier. Available at: http://linkinghub.elsevier.com/retrieve/pii/S0079612305510112 [Accessed January 21, 2016].

Merigan WH, Maunsell JHR (1993) How Parallel are the Primate Visual Pathways? Annu Rev Neurosci 16:369–402.

Merkulyeva NS (2022) Conducting Channels in the Visual System. The Third Channel. Neurosci Behav Physi 52:886–898.

Murray KD, Rubin CM, Jones EG, Chalupa LM (2008) Molecular Correlates of Laminar Differences in the Macaque Dorsal Lateral Geniculate Nucleus. J Neurosci 28:12010–12022.

Petry HM, Bickford ME (2019) The Second Visual System of The Tree Shrew. J of Comparative Neurology 527:679–693.

Ramón Y Cajal S (1909) Histologie du système nerveux de l’homme & des vertébrés. Paris: Maloine. Available at: https://www.biodiversitylibrary.org/bibliography/48637 [Accessed July 23, 2026].

Rathbun DL, Usrey WM (2009) Geniculo-Striate Pathway. In: Encyclopedia of Neuroscience (Binder MD, Hirokawa N, Windhorst U, eds), pp 1707–1710. Berlin, Heidelberg: Springer Berlin Heidelberg. Available at: https://link.springer.com/10.1007/978-3-540-29678-2_1976 [Accessed July 23, 2026].

Savier E, Eglen SJ, Bathélémy A, Perraut M, Pfrieger FW, Lemke G, Reber M (2017) A molecular mechanism for the topographic alignment of convergent neural maps. eLife 6:e20470.

Sciaccotta F, Kipcak A, Erisir A (2025) Morphological and Molecular Distinctions of Parallel Processing Streams Reveal Two Koniocellular Pathways in the Tree Shrew DLGN. eNeuro 12:ENEURO.0522-24.2025.

Shapley R, Hugh Perry V (1986) Cat and monkey retinal ganglion cells and their visual functional roles. Trends in Neurosciences 9:229–235.

Soares JGM, Botelho EP, Gattass R (2001) Distribution of calbindin, parvalbumin and calretinin in the lateral geniculate nucleus and superior colliculus in Cebus apella monkeys. Journal of Chemical Neuroanatomy 22:139–146.

Sugita S, Ho A, Südhof TC (2002) NECABs: a family of neuronal Ca(2+)-binding proteins with an unusual domain structure and a restricted expression pattern. Neuroscience 112:51–63.

Tasic B et al. (2018) Shared and distinct transcriptomic cell types across neocortical areas. Nature 563:72–78.

Tosches MA (2021) From Cell Types to an Integrated Understanding of Brain Evolution: The Case of the Cerebral Cortex. Annu Rev Cell Dev Biol 37:495–517.

Werneburg S, Fuchs HLS, Albers I, Burkhardt H, Gudi V, Skripuletz T, Stangel M, Gerardy-Schahn R, Hildebrandt H (2017) Polysialylation at Early Stages of Oligodendrocyte Differentiation Promotes Myelin Repair. J Neurosci 37:8131–8141.

Winer JA, Larue DT (1996) Evolution of GABAergic circuitry in the mammalian medial geniculate body. Proc Natl Acad Sci USA 93:3083–3087.

Yan Y-H, Winarto A, Mansjoer I, Hendrickson A (1996) Parvalbumin, calbindin, and calretinin mark distinct pathways during development of monkey dorsal lateral geniculate nucleus. J Neurobiol 31:189–209.

Yang H, Jeon C (1998) Distribution of calretinin in the superficial layers of the mouse superior colliculus: Effect of monocular enucleation. Korean Journal of Biological Sciences 2:389–393.

Zhang R, Long J-L, Ye Y-F, Ye H-Y, Zhao X-N, Cai X, Lu L (2025) Distributions of parvalbumin, calbindin-D28k, and calretinin in the cerebrum of Chinese tree shrews (Tupaia belangeri chinensis): A high-resolution neuroanatomical resource. Zool Res 46:893–911.

